# Assessing *in vivo* mutation frequencies and creating a high-resolution genome-wide map of fitness costs of Hepatitis C virus

**DOI:** 10.1101/2020.10.01.323253

**Authors:** Kaho H. Tisthammer, Caroline Solis, Faye Oracles, Madu Nzerem, Ryan Winstead, Weiyan Dong, Jeffrey B. Joy, Pleuni S. Pennings

## Abstract

Like many viruses, Hepatitis C Virus (HCV) has a high nutation rate, which helps the virus adapt quickly, but mutations come with fitness costs. Fitness costs can be studied by different approaches, such as experimental or frequency-based approaches. The frequency-based approach is particularly useful to estimate *in vivo* fitness costs, but this approach works best with deep sequencing data from many hosts, are. In this study, we applied the frequency-based approach to a large dataset of 195 patients and estimated the fitness costs of mutations at 7957 sites along the HCV genome. We used beta regression and random forest models to better understand how different factors influenced fitness costs. Our results revealed that costs of nonsynonymous mutations were three times higher than those of synonymous mutations, and mutations at nucleotides A or T had higher costs than those at C or G. Genome location had a modest effect, with lower costs for mutations in HVR1 and higher costs for mutations in Core and NS5B. Resistance mutations were, on average, costlier than other mutations. Our results show that *in vivo* fitness costs of mutations can be virus specific, reinforcing the utility of constructing *in vivo* fitness cost maps of viral genomes.

**Author Summary:** Understanding how viruses evolve within patients is important for combatting viral diseases, yet studying viruses within patients is difficult. Laboratory experiments are often used to understand the evolution of viruses, in place of assessing the evolution in natural populations (patients), but the dynamics will be different. In this study, we aimed to understand the within-patient evolution of Hepatitis C virus, which is an RNA virus that replicates and mutates extremely quickly, by taking advantage of high-throughput next generation sequencing. Here, we describe the evolutionary patterns of Hepatitis C virus from 195 patients: We analyzed mutation frequencies and estimated how costly each mutation was. We also assessed what factors made a mutation more costly, including the costs associated with drug resistance mutations. We were able to create a genome-wide fitness map of within-patient mutations in Hepatitis C virus which proves that, with technological advances, we can deepen our understanding of within-patient viral evolution, which can contribute to develop better treatments and vaccines.

## Introduction

Understanding the fitness costs of mutations in pathogenic viruses is critically important to improve strategies to prevent viral adaptation to host immune systems or to antiviral therapies [1–3], and for effective vaccine design. Viruses also provide an opportunity to study entire genomes to test hypotheses about the fitness costs of various types of mutations, allowing us to learn more about the process of evolution more generally. However, most studies on the fitness costs of mutations in viruses have been conducted *in vitro* [4–7]. Although studying viruses *in vitro* is often more practical than studying them *in vivo*, especially for viruses that infect humans, the results obtained may not reflect fitness costs in the real world. For example, viruses studied *in vitro* do not have to contend with the strong, multifarious selection imposed by a host’s immune response, resulting in a fitness landscape that may differ from the *in vivo* landscape in dramatic and important ways. Selective bias can also exist with culturing viruses itself [8]. In this study, we overcame the usual barriers to *in vivo* studies by analyzing clinical data obtained from 195 patients infected with the hepatitis C virus (HCV). We used these *in vivo* data to create a fitness cost map of the entire HCV genome.

To determine the *in vivo* fitness costs of mutations across the genome, we used information about (1) the HCV mutation rate and (2) a given mutation’s average frequency across multiple populations (*i*.*e*., patient samples) to determine the selection pressure against a given mutation at a given site. This ‘frequency-based’ approach [9] is based on a model of mutation-selection balance. As an RNA virus, HCV is especially well suited for this approach, and thus for testing hypotheses about how selection operates on the genome. Each patient infected with a long-lasting RNA virus infection harbors a diverse viral population that is almost perfectly independent of those in other patients [10]; thus, each patient sample in our study could be treated as a different viral population, as required in the frequency-based approach [9,11]. In addition, RNA viruses have mutation rates three to four orders of magnitude higher than those of bacteria and eukaryotes, due to the lack of proofreading activity in viral RNA-dependent RNA polymerases [12]. This allowed us to gather a great deal of information on fitness costs from a relatively compact genome.

The frequency-based approach has successfully been applied to estimate the *in vivo* fitness costs in RNA viruses: Theys et al. [9] showed fitness costs are higher for CpG creating mutations in the *pol* gene of HIV using data from 160 patients. Zanini et al. [11] reported genome wide *in vivo* fitness costs of mutations in HIV from nine patients, showing consistent estimated costs across patients, further validating the approach. However, the sample sizes were limited in both studies. In Theys et al. [9], the number of patients was relatively large (160) but the number of viral sequences were limited to about 20 per patient, totaling approximately 3000 data points per site. Deep-sequencing was used in Zanini et al. [11] with coverage per site in the order of thousands and higher, but the number of patients were limited to nine, totaling in the order of tens of thousands of data points per site. Using NGS data from a large number (195) of patients, our study overcame the sample size limitation by providing close to one million datapoints per site.

Estimating site-level fitness costs using the frequency-based approach requires mutation rates for each base substitution. The overall HCV mutation rate *in vivo* has been estimated to be around 10^−4^ to 10^−5^ base substitutions per site per replication cycle [7,13,14], though this was inferred from limited genome regions, and nucleotide-level substitution rates were not available. To overcome this limitation, Geller et al. [15] estimated the genome-wide HCV mutation rate using ultradeep sequencing *in vitro*, and found the overall rate to be 3.5 × 10^−5^ nucleotide mutation per replication cycle. Although the estimation was done on a modified replicon system with fragmented cistrons based on three cell lines, the rate estimated from their study was relatively consistent with previous *in vivo* estimates. Their ultradeep sequencing approach also allowed calculation of nucleotide-level mutation rates. Since no other base substitution rates are available for HCV, we applied the *in vitro* mutation rates from Geller et al. [15] to infer fitness costs.

By elucidating the *in vivo* fitness costs of point mutations in the coding regions of the HCV genome, we aimed to test several important hypotheses about fitness costs in general, as well as in HCV specifically. Using the HCV subtype 1a, first, we tested whether CpG-creating mutations are costly in HCV, as previous studies have shown that CpG usage tends to be under-represented in RNA viruses [16,17] and are highly costly in HIV [9]. Second, we determined how costly mutations are that result in drastic amino acid changes (*i*.*e*., the mutations that result in amino acid changes from one major group to another [positively charged, negatively charged, uncharged, hydrophobic and special cases]). Third, we assessed whether mutations that result in drug resistance are costly, and how prevalent the known resistance-associated variants are in pre-treatment patients.

## Results

### 1. Mutation frequencies within the HCV genome

First, we assessed *in vivo* mutation frequencies along the HCV1a genome using next-generation sequencing of samples from 195 HCV1a infected patients. We started by calculating genome-wide averages of mutation frequencies. The overall average genome-wide mutation frequency was 5.8 (± 1.3) × 10^−3^ per nucleotide, and transition mutations were around five times more common than transversion mutations (4.8 × 10^−3^ *vs*. 9.7 × 10^−4^) (Fig 1A and Table 1). For both transitions and transversions, the average frequency of synonymous mutations (transition: 8.1 × 10^−3^, transversion: 8.9 × 10^−4^) was more than twice as high than that of nonsynonymous mutations (transition: 3.3 × 10^−3^, Mann-Whitney test, corrected *P* = 0, transversion: 4.1 × 10^−4^, Mann-Whitney test, corrected *P* = 4.7 × 10^−175^). For transitions, the average frequency of nonsense mutations (2.0 × 10^−3^) was significantly lower than that of nonsynonymous mutations (Mann-Whitney test, corrected *P* = 3.9 × 10^−41^), but this was not the case for transversions. This suggests that the frequencies of nonsynonymous (4.1 × 10^−4^) and nonsense (5.2 × 10^−4^) transversions (*i*.*e*., approximately 4∼5 × 10^−4^) may reflect the lowest detection limit of our dataset (see Methods), since nonsense mutations are expected to occur at a lower frequency than nonsynonymous mutations. Since transitions mutations are observed at higher frequencies than transversions (likely because of their higher mutation rate), they offer greater statistical power and we decided to focus on transition mutations in all our subsequent analysis.

**Fig. 1.**
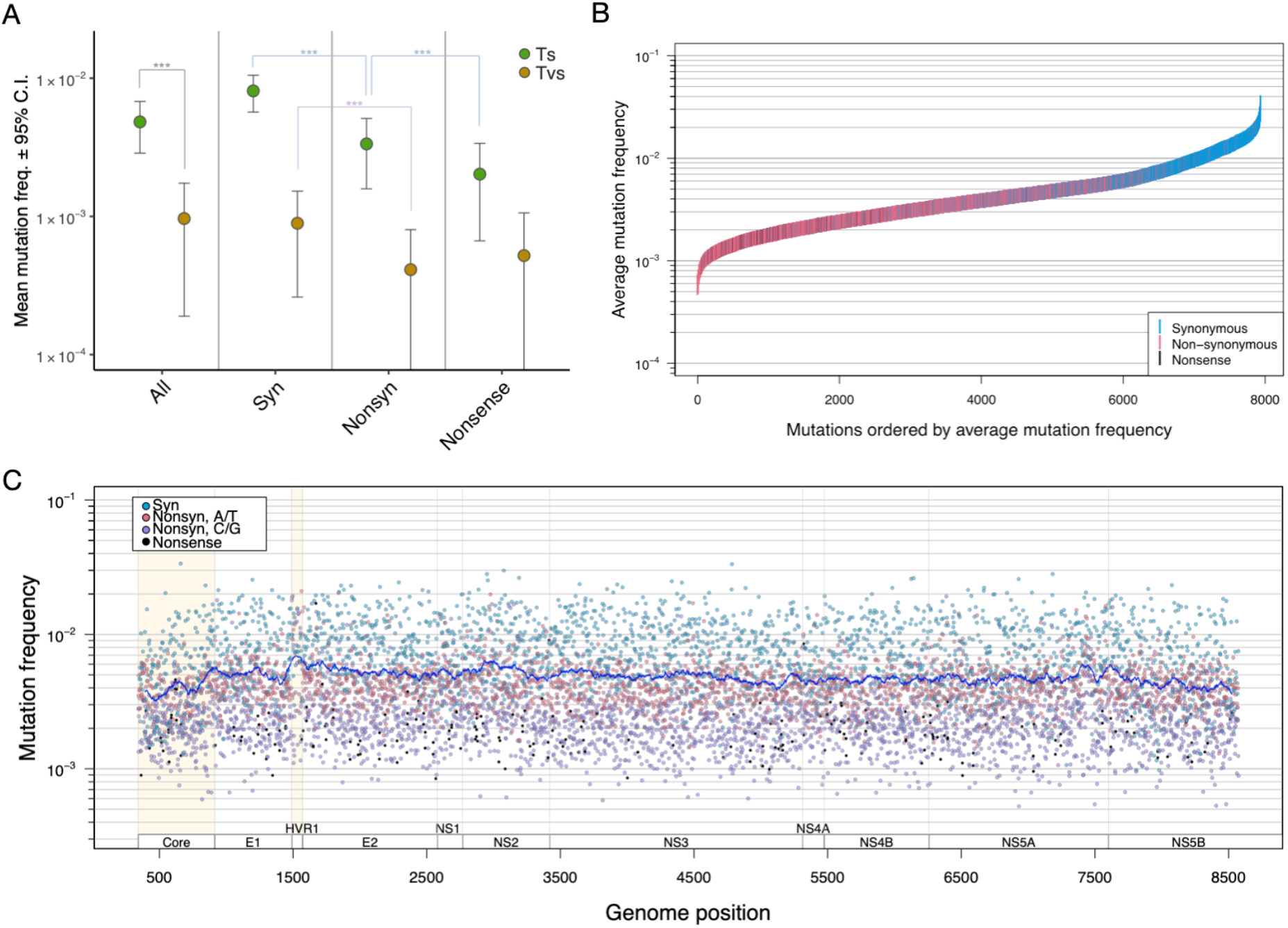
Estimated *in vivo* mutation frequencies in HCV1a, as determined by analysis of 195 viral populations. (A) Average estimated frequencies for all, synonymous (Syn), non-synonymous (Nonsyn), and nonsense mutations, stratified by transitions (Ts) *vs*. transversions (Tvs). Transition mutation frequencies are much higher for every class of mutation. ^***^ denotes statistical significance with adjusted P-values <0.001. (B) Genome-wide transition mutation frequencies, ordered by mutation frequency and colored by mutation type, show that synonymous mutations are more common than non-synonymous mutations in the HCV genome. (C) Transition mutation frequencies along the HCV genome, colored by mutation type, show that average mutation frequency is roughly consistent across the genome, with synonymous mutations more common than nonsynonymous mutations. Each dot represents the average mutation frequency at a nucleotide position, across the 195 viral populations. The line represents the sliding window average of 100 bases.

**Table 1.** Genome-wide average estimated frequencies of different types of mutation in the HCV1a genome.

| Mutation type | Average | 95% C.I. |
| --- | --- | --- |
| Transition | $4.8 \times 10^{-3}$ | $\pm 1.9 \times 10^{-3}$ |
| Synonymous | $8.1 \times 10^{-3}$ | $\pm 2.4 \times 10^{-3}$ |
| Nonsynonymous | $3.3 \times 10^{-3}$ | $\pm 1.7 \times 10^{-3}$ |
| Nonsense | $2.0 \times 10^{-3}$ | $\pm 1.4 \times 10^{-3}$ |
| Transversion | $9.7 \times 10^{-4}$ | $\pm 4.4 \times 10^{-4}$ |
| Synonymous | $8.9 \times 10^{-4}$ | $\pm 6.3 \times 10^{-4}$ |
| Nonsynonymous | $4.1 \times 10^{-4}$ | $\pm 3.9 \times 10^{-4}$ |
| Nonsense | $5.2 \times 10^{-4}$ | $\pm 5.4 \times 10^{-4}$ |

C.I. = Estimated confidence interval.

Next, we looked for trends in transition mutation frequencies across the genome. As expected, synonymous mutations generally had higher frequencies than nonsynonymous mutations. Nonsense mutations on the other hand were not clearly separated from the nonsynonymous mutations (Fig 1B). Throughout the genome, mutations at nucleotides A/T (*i*.*e*. A→G / T→C) had higher frequencies on average than those at C/G (*i*.*e*. C→T / G→A) (Fig 1C). The Hypervariable Region 1 (HVR1), the most variable section of the HCV genome [18,19], had the highest estimated average mutation frequency of all HCV genes (7.52 × 10^−3^, Mann-Whitney test (against each gene), corrected *P* = 1.1 × 10^−5^ ∼0.014, indicated in light yellow in Fig 1C, Table), whereas the Core gene, the genome’s most conserved gene [18,19], had the lowest average mutation frequency (4.07 × 10^−3^, statistically significant in 7 out of 10 genes, Mann-Whitney test, corrected *P* = 1.1 × 10^−10^ ∼ 9.3 × 10^−4^, indicated in light yellow in Fig 1C, also see Fig 2). The high average mutation frequency of the HVR1 region is due to higher nonsynonymous mutation frequencies than in other genes. Also, HVR1 was the only genic region with nonsynonymous mutation frequencies as high as synonymous frequencies (Fig 2).

**Fig. 2.**
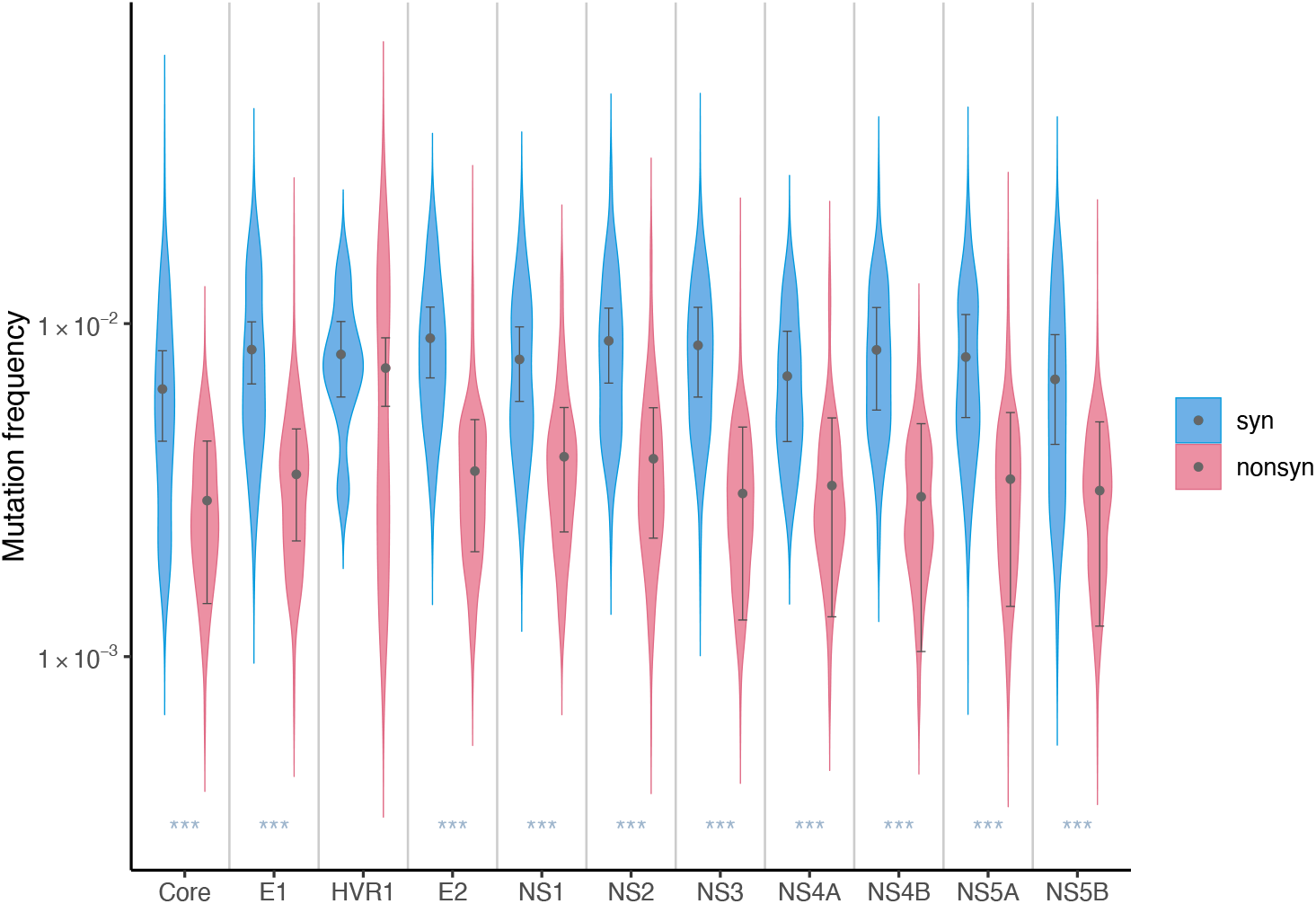
Estimated transition mutation frequencies of HCV by gene. Aggregated observed frequencies by gene and type of mutations; dots indicate the averages and the error bars represent the estimated 95% C.I. from 195 samples. ^***^ denotes statistically significant difference (adjusted P<0.0001) between synonymous and nonsynonymous mutations by Mann-Whitney test.

#### 1.1. Beta regression and random forest models show large effects of nonsynonymous mutations, nucleotide types, and genome location on mutation frequencies in HCV

In order to simultaneously assess the effects of multiple factors at the same time on HCV mutation frequencies, such as mutations that create drastic amino acid change or CpG sites, we used 1) a beta regression models since parameters of a beta regression are interpretable and they allow us to quantitatively assess the effects of specific factors, and 2) a random forest regression since the predictive machine learning (ML) approach allows us to investigate much larger numbers of factors (i.e. ‘features’ in ML) simultaneously and a random forest can give the relative importance of assessed features (but not the direction of the effects). A total of 83 features were include in our random forest regression analysis (S2 Table).

For the beta regression, we specifically explored the effects of the following factors; ancestral nucleotide, mutation type (synonymous/nonsynonymous mutations, CpG-creating mutations, mutations that result in a drastic amino acid change), genomic location (*i*.*e*., genes), RNA structure, and the interactions between the ancestral nucleotides and the synonymous or nonsynonymous mutation type on transition mutation frequencies. The results of the best-fit model, based on AIC, contained 18 variables and are shown in Fig 3A and S3 Table. Of the 18 variables, 17 had significant effects on mutation frequencies. The ancestral nucleotide had a large effect on mutation frequencies, with G→A and C→T mutations having 41∼51 % lower frequencies than A→G and T→C mutations (*P* < 0.0001) (Fig 3B). Nonsynonymous changes were also associated with a large reduction in mutation frequency of over 53% (*P* <0.0001) relative to synonymous changes. Our model also suggested relatively modest effects of genomic location when assessed by gene; the best-fit model included eight genes out of 11, of which seven had significant effects. The largest locational effect was with the HVR1 region, which had a frequency increase of 27 % (*P* <0.0001), whereas being located in Core was associated with a frequency reduction of 20% (*P* <0.0001). CpG-creating mutations and mutations that cause a drastic amino acid change were associated with modest reductions in mutation frequency of 7% (*P* < 0.0001) and 12% (*P* < 0.0001), respectively. Mauger et al. [20] identified 15 regions that likely form secondary structures and are functionally conserved in HCV genotype 1a, 1b, and 2a. We tested the aggregate effects of the 15 regions and found no significant effect on *in vivo* mutation frequencies. RNA structure was associated with an increase of ∼1.1% in mutation frequency (*P* = 0.39) and was not selected in the best fit-model. The interactions assessed showed modest effects sizes (−17 – 9.7%).

**Fig. 3.**
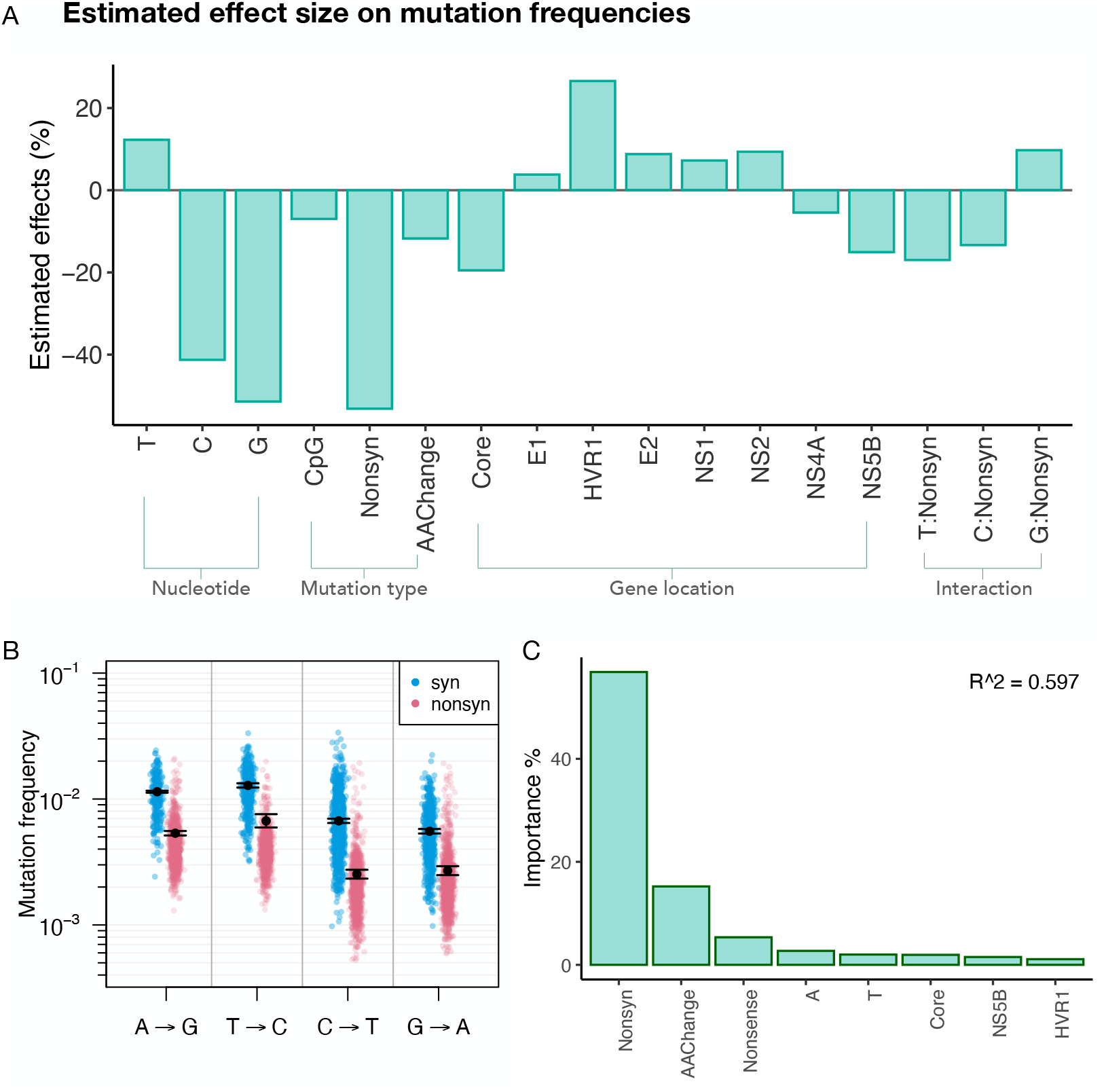
Various factors that affect the frequency of mutations in HCV, based on analysis of 195 viral populations, each derived from a different patient. (A) Predicted effects of ancestral nucleotide (T, C, or G, *vs*. A); CpG-creating status (*vs*. non-CpG-creating status); nonsynonymous (Nonsyn; *vs*. synonymous); amino acid-changing (AAChange; *vs*. non-amino acid-changing); presence in the Core, HVR1, E2, NS1, NS2, NS4A, and NS5B genes (*vs*. the NS3/NS4B/NS5A regions) on mutation frequencies in the HCV genome. Beta regression models were used to determine the effects of the different factors on mutation frequencies across the genome, and this figure reflects the results of the best-fit model based on AIC. (B) Estimated average transition mutation frequencies from the beta regression model (black dots with standard errors) and the actual observed frequencies from 195 patients infected with HCV (in colors). (C) Top 8 important features identified from the predictive random forest regression model on mutation frequencies (for all features tested, see S1 Table).

The results of the random forest regression model were highly congruent with the beta regression results, even though the predictability of our model reached a modest R^2 score of 0.597. The top features identified from the model were, in the order of importance, nonsynonymous changes, drastic amino acid changes, nonsense mutations (stop codons), ancestral nucleotide A and T, gene Core, NS5B and HVR1 in the order (Fig 3C, S1 Fig).

#### 1.2. *In vivo vs. in vitro* comparison reveals similar trends at the nucleotide-level, but different genome-wide patterns

We compared observed *in vivo* mutation frequencies with a study of *in vitro* mutation frequencies [15] (for transition mutations). Patterns of *in vivo* mutation frequencies were congruent with the trends observed in *in vitro* mutation frequencies [15] at the nucleotide level, but differed at the gene level. The *in vivo* estimated mutation frequencies were an order of magnitude higher than those found *in vitro*. Mutation frequencies for A→G and T→C mutations were higher both *in vivo* and *in vitro* than C→T and G→A mutations (S2A Fig). At the gene level, genome location was found to have very little effect (0.5%) on *in vitro* mutation frequencies [15], while our regression model results suggest that seven out of 11 genes have significant effects on *in vivo* mutation frequencies (Fig 3B and S1 Table). The difference was most prominent in HVR1; the mutation frequency peak observed in HVR1 *in vivo* was not seen *in vitro* (S2B Fig). The *in vitro* study [15] also found clusters of six highly mutable and 13 less mutable sites along the HCV genome, but in our *in vivo* study, we found no clear trends in mutation frequencies at the corresponding sites (S3 Fig). Similarly, the *in vitro* study [15] reported sites that had higher transversion than transition mutation frequencies (reversal sites) at over 1000 sites (12.4% of the total sites), but far less such reversal sites were observed *in vivo* (206 sites, 2.6%). The most salient reversal site *in vitro* (genome position C4080) again did not show a higher frequency of transversion mutations than transition mutations *in vivo*.

### 2. Genome wide fitness landscape of HCV

#### 2.1 Estimated selection coefficients reveal high costs of A→G and T →C mutations, nonsynonymous mutations, and mutations in NS5B

Based on the observed mutation frequencies, we next estimated selection coefficients (*s*) for each site in the HCV1a genome. Selection coefficients can quantitatively reveal fitness consequences of new mutations. We estimated *s* for each site based on the equation *f* = *µ*/*s* (see Methods), using the mutation frequencies (*f*) and the HCV mutation rates (*μ*) calculated from the datasets of Geller et al. [15]. The genome-wide average selection coefficient for transition mutations in coding regions, using the best available mutation rates [15], was 2.0 (± 0.7) × 10^−3^. The average selection coefficient for nonsynonymous mutations (2.5 ± 0.8 × 10^−3^) was three times higher than that for synonymous mutations (8.9 ± 4.8 × 10^−4^) (Fig 4A).

**Fig. 4.**
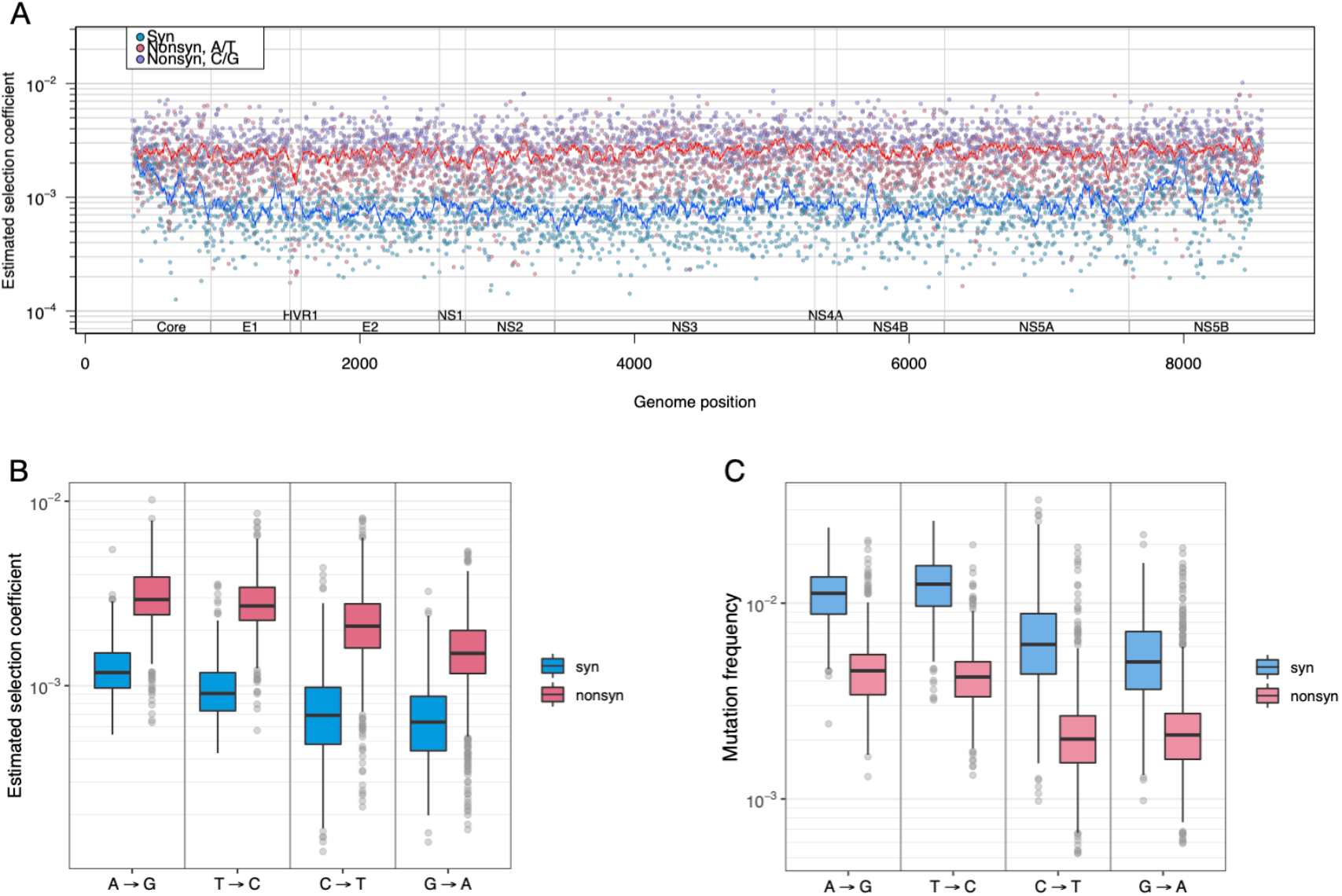
Estimated genome-wide selection coefficients (fitness costs) in the HCV genome. (A) Selection coefficients (1/replication cycle) along the HCV genome, colored by mutation type (syn = synonymous; nonsynon = nonsynonymous); each dot represents the average at each position across 195 patient samples, and lines represent the sliding window average for 50 bases. (B) Selection coefficients (1/replication cycle) stratified by nucleotide and syn/nonsyn status, colored by mutation type. (C) Estimated mutation frequencies stratified by starting nucleotide and syn/nonsyn status, colored by mutation type. Comparison of (B) and (C) shows higher estimated selection coefficients at A and T sites than at C and G sites, even though mutation frequencies were higher at A and T sites compared to C and G sites.

Selection coefficients for each nucleotide also revealed interesting patterns. The selection coefficients (*s*) and mutation frequency (*f*) generally have an inverse relationship (equation: *f* = *µ*/*s*); that is, higher *f* implies lower *s* (*i*.*e*., a lower cost). However, if mutation rates (*μ*) at different nucleotides vary widely, the inverse relationship may not hold when assessed at a nucleotide level, which was observed in our results. The average mutation frequency at nucleotides A and T sites (A→G and T →C mutations) was higher than at C and G sites (C →T and G →A mutations), leading one to expect that selection coefficients for A and T sites would be lower than those for C and G sites. In reality, however, for A and T sites, average selection coefficient values were higher despite the higher average mutation frequencies because the mutation rates based on Geller’s dataset [15] were so much higher (1.1-1.3 × 10^−5^ for A→G and T →C *vs*. 3.2 -4.2 × 10^−6^ for C →T and G →A) (Fig 4B and C). The differences in selection coefficients between nucleotides (excluding CpG-creating mutations) were all statistically significant, with the values of selection coefficients in descending order being A→G (1.5 × 10^−3^) > T→C (1.3 × 10^−3^) > G→A (8.4 × 10^−4^) > C→T (8.1 × 10^−4^) (Mann-Whitney test, corrected *P-* values = 0 – 3.5 × 10^−6^). It is, however, important to note that the degree of accuracy in estimating fitness costs (*s*) inevitably depends on the accuracy of estimated mutation rates. Therefore, we confirmed the that A→G and T →C mutations have a higher mutation rate than C →T and G →A mutations in the Geller data through a statistical test (Mann-Whitney test, all corrected *P*-values = 0.0), as well as calculating the 95% confidence intervals (S4 Fig).

Similarly, when comparing average selection coefficients for sites in different genes, the average selection coefficient for sites in NS5B (RNA dependent RNA polymerase) was the highest among all genes (statistically significant in 7 out of 9 genes, S5A Fig, Mann-Whitney test, corrected *P*-values = 1.2 × 10^−11^ - 0.077), despite the fact that Core’s average mutation frequency was the lowest among all genes (Table 2). This finding may reflect the higher genomic AT content of NS5B relative to Core (S5 Fig), since higher AT content can lead to a higher average selection coefficient due to A→G and T→C mutations having higher selection coefficients. Therefore, we also quantified the effects of different factors on selection coefficients using the two statistical models used for mutation frequencies: a beta regression and a random forest model.

**Table 2.** Average estimated selection coefficients per genic region of the HCV1a genome.

| Gene | s | 95% C.I. |
| --- | --- | --- |
| Core | $2.11 \times 10^{-3}$ | $\pm 1.1 \times 10^{-3}$ |
| E1 | $1.85 \times 10^{-3}$ | $\pm 8.2 \times 10^{-4}$ |
| HVR1 | $1.39 \times 10^{-3}$ | $\pm 1.0 \times 10^{-3}$ |
| E2 | $1.85 \times 10^{-3}$ | $\pm 9.0 \times 10^{-4}$ |
| NS1 | $1.66 \times 10^{-3}$ | $\pm 9.8 \times 10^{-4}$ |
| NS2 | $1.78 \times 10^{-3}$ | $\pm 1.1 \times 10^{-3}$ |
| NS3 | $2.03 \times 10^{-3}$ | $\pm 1.5 \times 10^{-3}$ |
| NS4A | $2.00 \times 10^{-3}$ | $\pm 1.6 \times 10^{-3}$ |
| NS4B | $2.01 \times 10^{-3}$ | $\pm 1.8 \times 10^{-3}$ |
| NS5A | $1.97 \times 10^{-3}$ | $\pm 1.7 \times 10^{-3}$ |
| NS5B | $2.21 \times 10^{-3}$ | $\pm 1.7 \times 10^{-3}$ |

S = Selection coefficient; C.I. = Estimated confidence interval.

#### 2.2. Beta regression and random forest models reveal nonsynonymous mutations, drastic amino acid changes, ancestral nucleotides, and key gene regions as determinant factors of fitness costs

We first applied a beta regression in a similar manner as to mutation frequencies, which followed highly similar trends as the mutation frequency results, except for the effects of ancestral nucleotide C and G for the reason described in the previous paragraph (Fig 5A, S4 Table). Model results also indicated that the effects of Core alone on selection coefficients was slightly larger than those of NS5B (+6.1% *vs*. +5.7%), likely reflecting the effects of higher AT content. As expected, sites in HVR1, which had the highest *in vivo* mutation frequencies, had the largest effect (−36.7%) out of all genes.

**Fig. 5.**
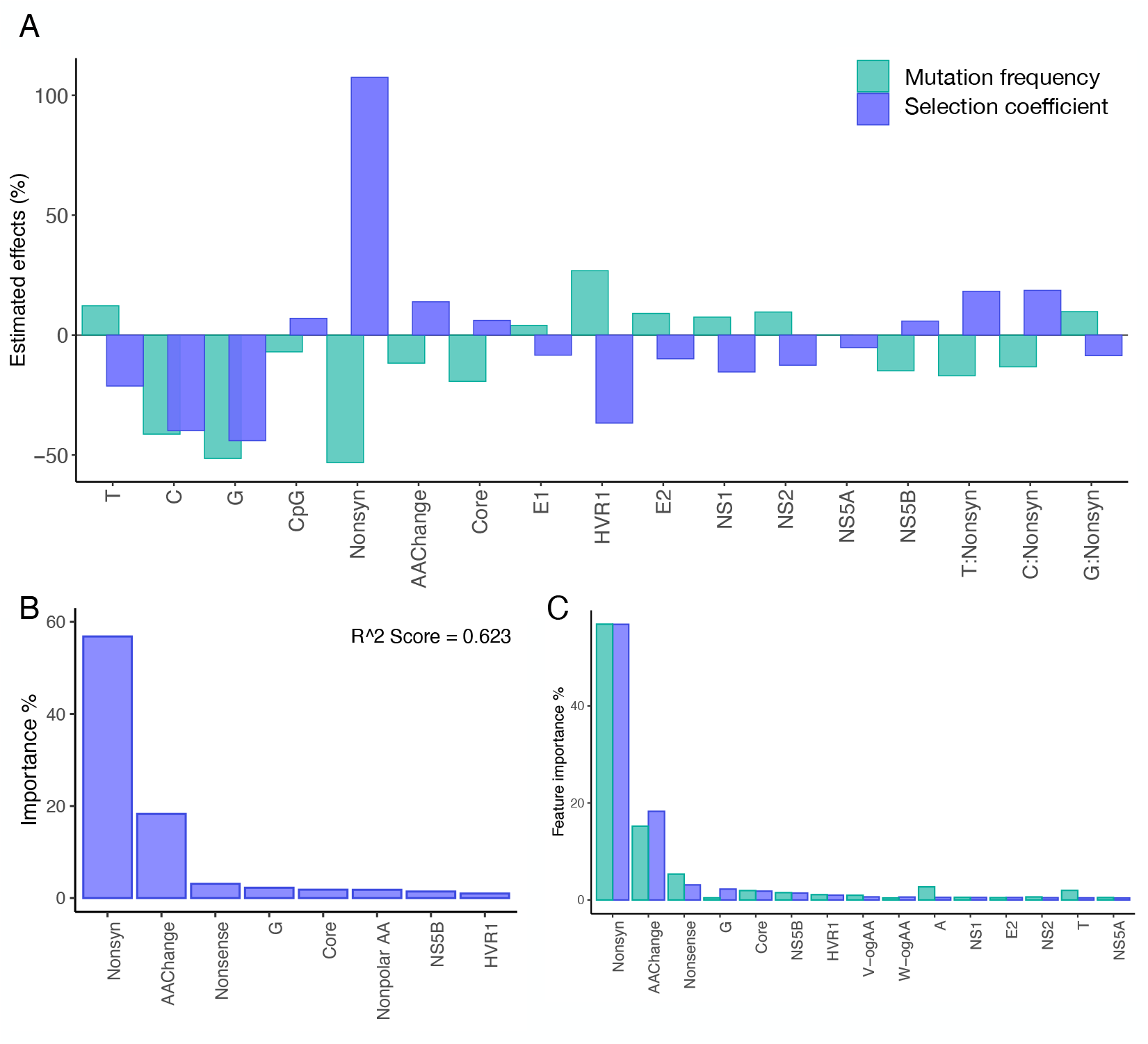
Factors that affect the fitness cost (selection coefficient) in HCV, based on analysis of 195 viral populations, each derived from a different patient. (A) Predicted effects from beta regression models on selection coefficients (SC) in the HCV genome, shown together with predicted effects on mutation frequencies (See Fig. 3A for description of different factors in the figure.) (B) Top 8 important features identified from the predictive random forest regression model (for all features tested, see S1 Table). (C) Overlapping features ranked in the top 20 in the random forest regression models for mutation frequencies (MF) and selection coefficients (SC).

We also applied the RF regression models to further elucidate contributing factors for selection coefficients in the same manner as for mutation frequencies. Features identified as important for predicting selection coefficient values were in agreement with our beta regression outcomes. The results identified ‘nonsynonymous mutation’ as the most determinant factor (Fig. 5B), followed by drastic amino acid changes, nonsense mutations, ancestral nucleotide G. Gene locations of Core, NS5B and HVR1 were also in the top eight features of importance. We then compared the features that ranked top 20 in both RF models for mutation frequency and selection coefficients, which showed highly consistent results, revealing the three most important determinants as nonsynonymous mutations, drastic amino acid changes and nonsense mutations in both models (Fig 5C).

#### 2.3. Distribution of fitness effects and gamma parameter estimation

Based on the estimated fitness costs (*s*) for each nucleotide in our study, we assessed the distribution of fitness effects (DFEs) of synonymous and nonsynonymous transition mutation for each nucleotide (S6 Fig). All synonymous mutations and 99.7% of nonsynonymous mutations had fitness costs <0.01. Parameters for the gamma distribution were also estimated for the entire DFEs. The scale (k) and shape (θ) parameters were estimated to be 8.32 × 10^−4^ (95% confidence interval [CI] = 8.11 – 8.55 × 10^−4^) and 2.378 (95% CI = 2.318 -2.436), respectively, which can be applied to future population simulation studies of HCV.

#### 2.4. High fitness costs of drug resistance sites

A number of direct-acting antiviral agents have been developed for HCV, all of which specifically target the NS3, NS5A, or NS5B proteins. Drug resistance often develops in patients, and many HCV resistance-associated variants have been reported [21–26]. Prevalence of baseline (pre-existing) resistance-associated variants in treatment naive patients (*i*.*e*. natural occurrence), as well as the fitness costs of these variants provide insights into the evolution of drug resistance traits, treatment outcome prediction, and durability of resistance phenotypes [27–29]. They can also help guide treatment options, such as combining antivirals for which resistance mutations have high fitness costs to lower the risk of developing drug resistance.

We assessed the natural occurrence of resistance-associated variants using’our ‘unfil’ered’ HCV datasets, which included sites with alternate majority nucleotides and transversion mutations, as opposed to the filtered datasets used to estimate *in vivo* fitness costs, which excluded the sites that were not in selection-mutation balance, including those with a nucleotide different from the reference (H77, AF011753, see Methods). Of 87 known resistance-associated variants across 26 sites we examined (S5 Table), natural occurrence was observed for all variants in our datasets, including the 51 variants previously not observed or reported [23] (Fig 6). The proportion of the samples with pre-existing resistance-associated variants varied from 3.6 to 99.5% (average 62%) depending on the resistance mutation (S7 Fig.). The result highlights that many variants do occur naturally in treatment naive patients, making them more vulnerable to developing drug resistance. Additionally, 20 of the 87 resistance-associated variants were observed as a majority variant in one or more patients (S7 Fig, ‘% fixed sample’). The percentages of patients that had the resistant variant as the majority vary from 0.5 % to 54.6%, depending on the variants, with Q80K in NS3 being most likely to be present as the majority. The average fitness costs of all resistance-associated variants examined was 4.7 × 10^−3^, with 2.5 × 10^−3^ for transition mutations and 6.8 × 10^−3^ for transversion mutations. Estimated fitness costs of resistance-associated variants (at transition sites only) were compared to those of non-resistance-associated sites in the three genes, which revealed on average 21% higher fitness cost in resistance-associated variants of HCV (Mann-Whitney test, *P* = 0.0021).

**Fig. 6.**
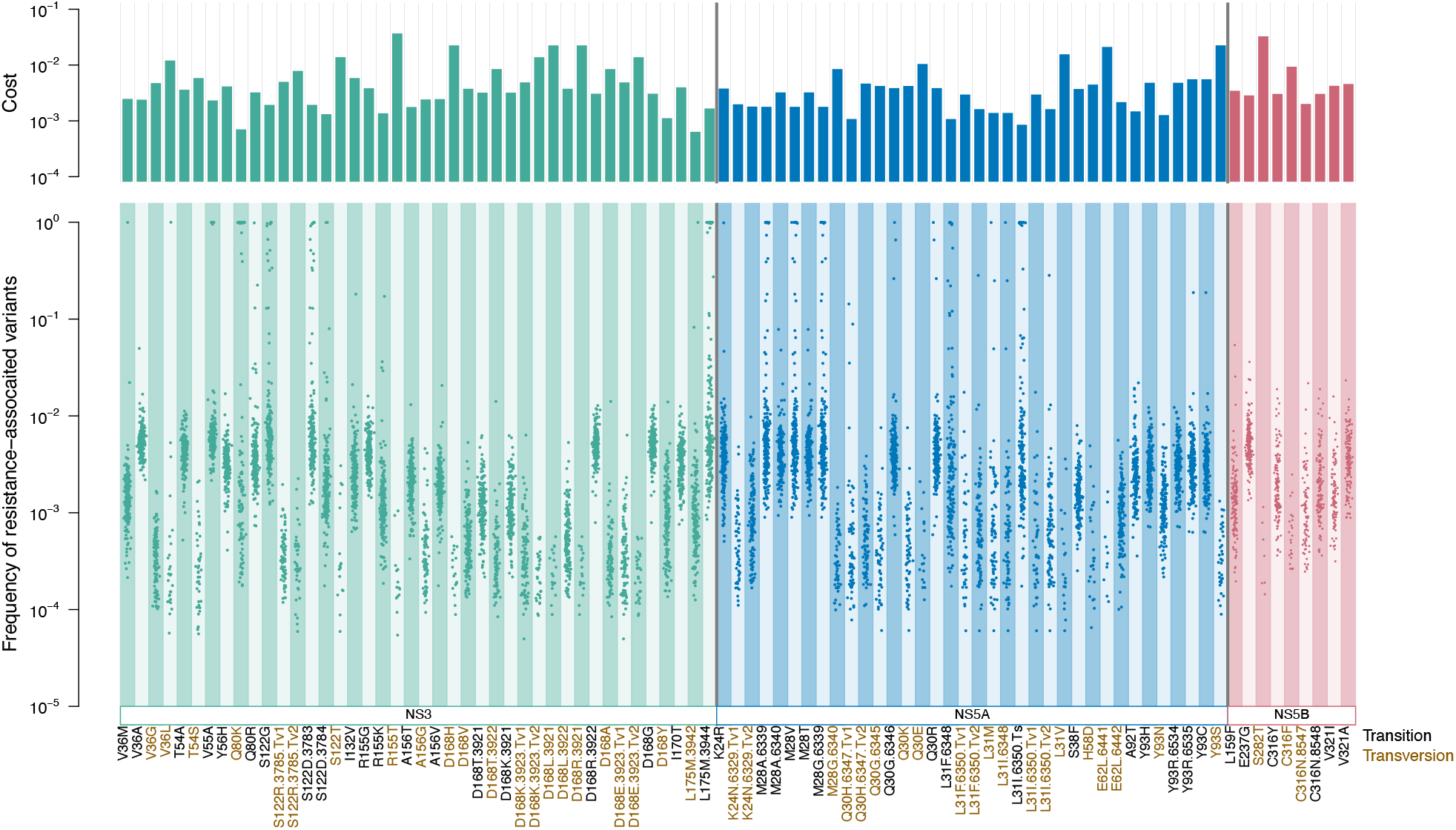
Occurrence and fitness costs of resistance-associated variants in the HCV genome. Estimated fitness costs (top) and natural prevalence (bottom) of resistance-associated variants, where each dot represents a mutation frequency observed in a patient. Variant names in black are created by transition and those in brown are created by transversion mutations. RAVs that can be created by different mutations are specified in names, in a format of variant name, nucleotide position, and type of mutation (Tv1 stands for transition mutations that result in C or A and Tv2 stands for transition mutations that results in G or T).

### 3. Validation of our frequency-based approach

The frequency-based approach we used in this study assumes that site-level mutation frequencies are governed by consistent mutation rates and selection coefficients (mutation-selection equilibrium) across different viral populations (*i*.*e*., patients). To confirm this assumption, we ran simulations to generate site-based mutation frequencies and compared these to observed frequencies. We also compared the patterns of *<u>in vitro</u>* mutation rates from the study of <u>[1]</u> with the patterns of *in vivo* mutation rates estimated in our study. Additional analyses that we performed to further to validate the frequent based approach are summarized in S2 Text (SI File).

#### 3.1. Simulated mutation frequencies based on the estimated selection coefficients are consistent with observed mutation frequency patterns

To confirm that the data are consistent with HCV populations being in mutation-selection balance, we created simulated datasets under the assumption of mutation-selection balance and found that simulated and observed frequency distributions were very similar. Specifically, we ran simulations to generate mutation frequencies using the nucleotide specific mutation rate estimates from <u>[1]</u>, selection coefficient estimates from our dataset for each site, and a within-host viral population size of 100,000 using code written previously <u>[1]</u>. Using these input parameters, for each site separately, we simulated 195 viral populations (one for each patient). In these simulated populations, we follow a population of 100,000 virus-infected cells. Each simulated generation, 100,000 new cells are infected using a random sample of virus from the previous generation. Mutations occur during replication, and virus with mutations has a slightly lower probability to infect new cells in the next generation (quantified by the site-specific fitness cost). We then “harvest” samples from these simulations of mutation-selection balance and determine how many cells carry mutated virus, to calculate the mutation frequency for that site for that patient. For each of the 7957 sites, we compare the simulated mutation frequencies with the observed mutation frequencies using a Mann-Whitney test. We found that for 92.6 % of the sites the distribution of frequencies was not significantly different than the simulated distribution of frequencies (*i*.*e*., adjusted P-values > 0.05 by Mann-Whitney Test with Holm correction). The remaining 7.4 % of sites showed significantly difference between observed and real data. This could be due to these sites not being in mutation-selection equilibrium, due noise in the NGS data which is not reflected in the simulated data, or due to epistasis or patient-specific fitness effects. This result demonstrates that most of the data we used in this paper are consistent with a model of mutation-selection-equilibrium (S8 Fig., S1 Datafile).

#### 3.2. Patterns of *in vitro* mutation rates correspond to those of *in vivo* estimates of mutation rates

The mutation rates used to obtain fitness costs in our study came from *in vitro* data [15], since *in vivo* nucleotide-level mutation rates are not currently available. If *in vivo* and *in vitro* mutation rates differ drastically, it would affect our estimations of fitness costs. Therefore, we evaluated pseudo *in vivo* mutation rates using nonsense mutation frequencies observed in our dataset, assuming that the fitness cost (*s*) of a nonsense mutation is one [6]. There are nine base substitution combinations (out of 12) that can create stop codons; specifically, since there are no Cs in stop codons, A→C, T→C, or G→C cannot be nonsense mutations. All but G→T nonsense mutations were observed in our datasets. The pseudo mutation rates obtained from our *in vivo* nonsense mutation frequencies followed a similar pattern as *in vitro* rates, with a significant correlation between the two groups (Spearman’s ρ = 0.79, *P* = 0.028) (S9 Fig).

## Discussion

Genome-wide maps of fitness costs at each locus are key to understanding the evolutionary dynamics of pathogenic viruses, yet few such maps exist. Here we used deep sequencing data from 195 viral populations (*i*.*e*., sequencing data from 195 infected hosts) and applied the frequency-based approach to estimate *in vivo* fitness costs. We present a high-resolution map of inferred *in vivo* fitness costs for transition mutations at 7957 sites in the HCV1a genome. Our results revealed that whether a mutation is nonsynonymous, the ancestral nucleotide, and genic location are the most important factors that determine fitness costs.

### Comparing to HIV

Previous studies on fitness costs of mutations in HIV found similar trends in site-level fitness costs, such as a clear difference in synonymous and nonsynonymous mutations and large effects of the ancestral nucleotide [9,11]. However, there are also clear differences between HIV and HCV. CpG-creating mutations and mutations that cause drastic amino acid changes have large effects on fitness costs in the HIV pol gene, have markedly smaller effects in HCV, suggesting that the magnitude of these effects likely vary between viruses. Specifically, CpG-creating mutations are 1.5 times more costly than non-CpG creating mutations in HIV, but only 1.07 times more costly than non-CpG creating mutations in HCV. Many RNA and DNA viruses display suppressed frequencies of CpG creating mutations, indicating that CpG creating mutations are costly to a variety of viruses [17], and the magnitude of such effects appear to vary greatly among viruses [16]. This may be due to different interactions of these viruses with our immune systems or differences in their replication cycles and merits further study.

### Comparing *in vivo* mutation frequencies to *in vitro* mutation frequencies

*In vitro* studies conducted in tightly controlled environments allow us to obtain detailed information about viral evolutionary dynamics, including nucleotide level mutation rates. However, the use of cell-lines in an environment without selective pressures from hosts creates different dynamics from *in vivo* evolution. Understanding the difference and similarity between *in vitro* and *in vivo* evolution can offer crucial insights into which forces are important for viral evolution *in vivo*. Here we compare the *in vivo* mutation frequencies from our study with *in vitro* mutation frequencies from the study of <u>[1]</u>.

The *in vivo* mutation patterns from our study highlighted the selective pressures exerted by the host immune system on different parts of the HCV genome, compared to those of *in vitro*. Significant location effects were observed *in vivo*, but not *in vitro* [15]. First, the prominent peak in mutation frequency in HVR1 was observed *in vivo*, while no such peak was observed *in vitro*. HVR1 plays an important role in immune escape through positive selective pressure [18,30], and we observed that HVR1 was the only genic region with similarly elevated nonsynonymous mutation frequency relative to synonymous frequency (Fig 2B). In addition to the effect of HVR1, we also observed location effects for seven other genic regions, which likely reflects selective pressures from the host, which are not present in the *in vitro* study. Lastly, the sites with high mutability identified by <u>[1]</u> *in vitro* did not show similar trends *in vivo*, suggesting that these sites are also under strong negative selection *in vivo*.

We found several important similarities between the *in vivo* and *in vitro* settings. There was no significant effects of RNA structure on either *in vivo* and *in vitro* mutation frequencies [15]. We found that mutation frequency patterns at the nucleotide-level appear to be consistent between *in vivo* and *in vitro*; A and T have higher mutation frequencies than G and C in both cases [15]. Mutation rates for each nucleotide were also correlated between *in vivo* and *in vitro* (*i*.*e*., relative rates of different nucleotides were consistent), where we used observed frequencies of nonsense mutations to estimate *in vivo* mutation rates (Fig S9). This is important since it validates our application of *in vitro* mutation rates to estimate *in vivo* fitness costs. However, the *in vivo* mutation rates were two orders of magnitude higher than those of *in vitro*. We suspect that this is partly because of the costs of *in vivo* nonsense mutations are not likely to be equal to one as we assume *in vivo*; defective viruses with premature stop codons can be rescued by other defective or non-defective viruses [14,32–35] due to complex quasispecies dynamics in viruses [36,37]. This phenomenon may allow nonsense mutations to be observed at higher-than-expected frequencies *in vivo* viral populations. Also, how we calculated nonsense mutation frequencies was not based on counting actual stop codons; we calculated the site-level nonsense mutation frequencies, which would create stop codons if other two nucleotides in a codon were the same as the reference. This would slightly overestimate the actual nonsense mutation frequencies. Ribeiro et al. [14] estimated the *in vivo* mutation rate using a model and validated with nonsense by counting all stop codons. Their estimation (2.5 × 10^−5^ mutations per nucleotide per replication cycle) aligned with the *in vitro* mutation rate estimation [15], which may be due to their PCR-based sequencing, which cannot capture low frequency mutations as NGS can and their stricter method of calculating stop codons (Text S1 in [14]).

### Prevalence of drug resistance

Natural occurrence of resistance associated variants was more frequently and widely observed in our study than previously reported [23]. This highlights the importance of deep sequencing, since many variants occur at a low frequency and would not have been captured by traditional Sanger based methods. Although the fraction of patients in which the resistance-associated variant was fixed was fairly low (average 5.7% of patients), one variant Q80K was observed at an especially high prevalence (fixed in 54.6% of patients, the variant was observed in 94.3% of patients, S7 Fig). This is consistent with previous observations, as Q80K is known to exist at a high prevalence (up to 48%) in strains from North America [21,23,38]. The results provide important insight into how common resistance associate variants are.

### Strengths and limitations of this study

Deleterious mutations that occur at a low frequency indicate high fitness costs. However, our results suggest that inferring nucleotide-level fitness costs from mutation frequencies should be done cautiously: Lower mutation frequencies may not necessarily translate into higher fitness costs, if substantial differences in mutation rates exist at the nucleotide level along with biased base composition. In our dataset, mutations at A or T sites had higher fitness costs than mutations at C or G sites, even though their mutation frequencies were significantly higher. A similar trend was observed in a previous study on HIV’s *pol* gene; *in vivo* mutation frequencies were highest for G→A mutations even though fitness costs were also highest for G→A mutations, due to the high mutation rate at these sites [9]. Thus, without nucleotide-level mutation rates, differences in fitness frequencies at the nucleotide-level need to be interpreted carefully.

Our unique approach worked well, thanks to the fact that the average read depth at each site was > 6,200 after filtering, which resulted in an average of > 877,000 data points at each nucleotide site analyzed. The correlation between *in vivo* and among-host genetic diversity at each site supports our basic assumption that *in vivo* mutation patterns are largely dictated by intrinsic factors at selection-mutation equilibrium. Adding > 400 genomes from NCBI had little effect on the correlation between *in vivo* and among-host diversity, which suggests that we had more than adequate sample size to capture the true genetic diversity of HCV1a. Additionally, we have previously performed simulation studies to assess the optimal combination of sequencing depth and sample size, which showed that > 150 samples with at least 1,000X sequencing depth should yield reliable estimates (*i*.*e*., a high correlation coefficient, > 0.95, between real and estimated fitness costs, S10 Fig). Here, we had 195 samples with 6200X sequencing depth, more than enough to produce reliable data.

## Conclusion

Despite HCV’s extremely high level of genetic variation, our detailed analysis of genome wide mutation patterns revealed *in vivo* mutations do not occur in a completely stochastic manner, illuminating the mutant spectrum complexity of HCV. Swarms of mutants, often termed viral quasispecies [36,37], are continuously generated in RNA virus populations. The mutation spectrum composition varies over time, producing transient equilibrium distributions that are affected by internal and external factors [32]. However, even with these ever-changing mutant spectra, our results showed that we could observe a situation close to mutation-selection equilibrium. RNA viruses are considered to replicate at a mutagenesis level close to the lethal threshold, at which the error rates exceed the tolerance limit of genetic information preservation [39]. A delicate balance between this threshold and host selective pressures is likely responsible for shaping the observed *in vivo* mutation patterns; approximately 50% of sites were highly conserved while the remaining variable sites contribute to continuous generation of variation. From a clinical perspective, one of the most challenging aspects of the treatment and management of pathogenic RNA viruses is their high genetic diversity. Targeting the highly conserved, high-cost sites or regions may help improve treatment efficacy by hampering the development of drug resistance.

This study serves as an additional proof of concept for the frequency-based approach to understanding *in vivo* fitness costs. This opens the door to opportunities to investigate *in vivo* fitness costs of a wide variety of rapidly evolving pathogens, as deep sequencing of pathogen populations becomes more common with the steep decline in next generation sequencing costs. We showed that there are aspects of viral evolution that we cannot infer from *in vitro* studies, as well as from the between-host level of analysis. Practical applications of better understanding *in vivo* evolution and the fitness landscape of viruses include more targeted antiviral drug development, efficient treatment management, better treatment outcome prediction, and informative for vaccine design. By leveraging the technological advances and tremendous resources already available, we have the potential to deepen our understanding of *in vivo* evolution of viruses.

## Methods

### Samples

We obtained stored baseline plasma samples from a selection of HCV treatment naive and viremic HCV/HIV co-infected participants from sites in the Canadian Co-infection Cohort (CCC) [40]. Samples were collected between May 8, 2003 and November 20, 2014 from acute or chronically infected patients attending 18 different clinics across Canada. We used 195 samples identified as genotype 1a out of the dataset for our study.

### Viral RNA extraction and sequencing

For each baseline sample HCV RNA was extracted from 500 ul of plasma using a NucliSens easyMag (BioMerieux Inc.). Extracted RNA was eluted in 60ul of elution buffer and stored at -20C prior to amplification. Near full length amplicons were generated using an oligo d(A)20 primer for cDNA synthesis. Two nested PCR reactions using pangenotypic primers were utilized to generate an 8,991-bp amplicon spanning the HCV Core to NS5B regions. Subsequent to second round PCR amplification MiSeq sequencing libraries were generated using Nextera XT DNA Sample Preparation Kits (Illumina, Inc.). Amplicons for each sample were tagged using Nextera XT Index Kits (Illumina, Inc.), multiplexed and sequenced on an Illumina MiSeq next-generation sequencing platform (Illumina, Inc.).

### Data filtering and formatting steps

Sequence reads of virus isolates were quality filtered and trimmed using BBTools v38.38.00, following its suggested steps [41]. First, the adapters were removed. To increase our confidence in identifying minor variants, bases that had phred score <35 were all trimmed from both ends. Contaminants (PhiX) were then filtered out and reads with a <35 average score were also eliminated. Potential duplicates were identified and clumped, and filtered reads were merged. To retain maximum read depths in order to confidently estimate minor variant frequencies, both merged and unmerged reads were mapped to a reference sequence using BWA 0.7.17 [42]. Mapped reads were hard clipped, and the resulting SAM files were imported into R and filtered further to eliminate potential contaminants and chimeric sequences. This was done using a custom script in which all reads with a harmonic distance >10 from their own consensus sequences were eliminated, as they likely represented contaminants or mis-mapped sequences. This cut off value was determined empirically; histograms of read counts per harmonic distance in each sample showed that the first peak mostly flattened out by 10, and the second peak, which represents mis-mapping, did not appear until past 10. Filtered reads were remapped to their own consensus sequences using BWA, and the resulting bam files were imported in R [43]. For each patient sample (*i*.*e*., viral population), frequency tables of nucleotides at each position were created with Rsamtools and custom scripts in R.

Each frequency table was processed to calculate the frequencies of transition and transversion mutations relative to the reference sequence (wildtype H77). Types of mutation (synonymous/nonsynonymous, CpG-creating, those causing drastic amino acid changes, etc.) were also identified for each site. To accurately estimate *in vivo* deleterious mutation frequencies at the selection-mutation-drift equilibrium, we applied rigorous filtering, removing sites with a mutation frequency >0.2 (*i*.*e*., those likely under active selection), sites that had a different majority nucleotide from the reference, and sites with <1000 read depth. Mutation frequencies of all samples were then compiled into one table, and sites that were present in less than one-third of the total sample (< 65) were removed. The filtering steps produced 7,958 usable sites in the coding regions, and the averages of mutation frequencies were calculated for these sites. This process yielded a table containing the average transition mutation frequencies at each position across the HCV genome. To assess the natural occurrence of RAVs, the frequency tables were processed similar to the previous step, but further filtering steps were not performed except for removing sites with <1000 read depth. The 95% confidence intervals for mutation frequencies/selection coefficients were estimated by calculating standard errors 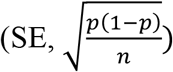 at each site for each patient based on the number of reads (n), and then means of SEs for each site were calculated across all patients to obtain the global estimates of standard errors at each site. Using these estimated standard errors, upper and lower 95% confidence intervals were estimated by z_ɑ=0.05_(=1.96) x SE for each site. We used this estimation approach as aggregating at a patient level was not optimal because of resulting in losing sample size and thus leading to overestimation of standard errors [44].

### Data validation

Since the sequencing error rate of the Illumina Miseq is reported to be around 0.1% [45], we took additional steps to validate our filtering methods and the resulting datasets. We were concerned that if our previous filtering steps did not remove the errors efficiently, mutations observed at low frequencies (<0.001) could actually represent sequencing errors. To evaluate whether this posed a problem, we replaced all mutation frequencies <0.001 with either NA or zero and assessed the resulting mutation frequency patterns. The results showed consistent patterns. Therefore, we used the original datasets for further analysis. Additionally, we have used the homogeneous sample of RNA virus (SIV) from a different study to estimate the error rates from our pipeline [46]. The results were consistent with published previous MiSeq error rates: The average mutation frequency of the ‘homogeneous viral sample’ was 0.00156, and its median was 0.00148. The error rate estimations confirmed that our approach can estimate mutation frequency lower than these values since our transversion mutation frequencies were 0.00089, clearly lower than 0.001, and biologically makes sense. We also assessed whether viral load biased observed mutation frequencies by comparing the average mutation frequencies of patient samples with low and high viral loads. We found no difference in the average mutation frequencies between the two groups. Therefore, all samples were included in the analysis.

### Calculation of selection coefficients and other statistics

To gain a fine-grained understanding of the *in vivo* fitness effects of mutations in HCV viruses, we used the *in vivo* frequency-based approach [9,11], in which fitness costs of mutations are estimated from their frequencies across multiple natural populations (*i*.*e*. multiple patients). This approach relies on translating *in vivo* mutation frequencies into fitness costs based on the principle of mutation-selection balance. In infinitely large populations, the opposing forces of mutation and selection cause costly mutations to be present at a constant frequency [47,48]. In haploid systems, it is expressed as:

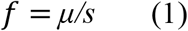

where *μ* is the mutation rate from wildtype to mutant and *s* is the negative fitness effect of the mutation (selection coefficient). In natural populations of finite size, the frequency of mutations is not constant but fluctuates around the expected frequency of *μ/s*, because of the stochastic nature of mutation and replication [49,32]. It is thus impossible to accurately infer the selection acting on individual mutations from a single observation of a single population. We solved this problem by averaging frequencies of each individual mutation from multiple populations (sampling vertically, rather than horizontally, S11 Fig.). Note that the principle of mutation-selection balance holds even when bottlenecks or selective sweeps occur, because such events do not change the expected frequencies of mutations. Also, this method allows us to directly assess fitness costs at all sites in the genome and to calculate selection coefficients rather than relative rates [9].

Selection coefficients were estimated for each mutation at each nucleotide based on the equation (1) using estimated *in vivo* mutation frequencies and mutation rates calculated from raw datasets of [15] in R, using the same method Geller et al. [15] used to calculate the global mutation rate. Briefly, mutation rates for each nucleotide-level mutation were obtained by dividing the total mutations observed during the experimental period by the generation cycle with adjustment for errors. The nucleotide-level mutation rates are the keys for estimating selection coefficients using the frequency-based approach. Therefore, we assessed the validity of *in vitro* mutation rates from Geller et al. [15] using beta regression (see below), and found no major effects of selective pressures seen in *in vivo* data (see S2 Text SI File). *In vitro* mutation frequencies used in comparison were also derived from the datasets of [15].

Statistical analyses were performed in R and Python. Beta regression model fitting was conducted using the R package *betareg* [50]: For predictor variables, we included ancestral nucleotide, types of mutations (synonymous vs. nonsynonymous, create a CpG site or not, cause a drastic amino acid change or not), location (11 genic regions), RNA structure (the aggregate effects of 15 regions that likely form secondary structure, identified by [20]), and interactions. AIC values were used as a criterion to select the best fit model. Parameters for DFEs were estimated using the R package noplr [51]. To assess the diversity between *in vivo* and among hosts, additional HCV1a genomes were downloaded from NCBI GenBank. Random forest regression analyses were conducted in Python 3.9 (Python Software Foundation, https://www.python.org/) using Scikit-learn [52] module. After constructing a matrix of all features to be used in RM (83 features including mutation types, nucleotides, amino acids, types of amino acids, location, and RNA structure, see S3 Table), highly correlated features were removed, and hyperparameter tuning was performed using the GridSearchCV and RandomizedSearchCV functions in Scikit-learn. Models fitted with the best hyperparameters reached R^2^ of 0.597 for mutation frequencies and 0.623 for selection coefficients.

Simulation of site-specific mutation frequencies were performed using the nucleotide specific mutation rates estimated from <u>[1</u>], site-specific selection coefficients estimated in our current study, and a population size of Ne = 100,000 to generate mutation frequencies for the total number of samples available for the particular site (ranged from 65 to 189). The simulation program is compiled in C++ and available at the GitHub repository, along with all coding scripts (https://github.com/kahot/HCV_FitnessCost). We then sampled from these simulated populations using on the average read depths of each patient. The resulting simulated sample frequencies and the observed frequencies were statistically compared using the Mann-Whitney test with the Holm correction.

## Acknowledgments

The authors thank Kristin Harper, PhD, of Harper Health & Science Communications, LLC, for providing writing and editorial support in accordance with Good Publication Practice (GPP3) guidelines.

## Supporting Information

**S1 Fig. Top 20 features identified as important from the random forest regression model for predicting *in vivo* HCV mutation frequencies**.

**S2 Fig. Comparison of *in vivo* and *in vitro* transition mutation frequencies in the HCV1a genome**.

**S3 Fig. Transition mutation frequencies of the sites corresponding to the highly mutable (HM) and less mutable (LM) sites identified in Geller et al. (2016)**.

**S4 Fig. Mutation rates estimated from the dataset obtained from Geller et al. (2016)**.

**S5 Fig. Average selection coefficients (A) and AT contents (B) of each genic region in the HCV genome**.

**S6 Fig. Distribution of fitness effects for each nucleotide (A, T, C, G) in the coding regions of the HCV genome, stratified by synonymous and nonsynonymous mutation status, using the estimated fitness costs (s) calculated in this study**.

**S7 Fig. The percentage of 195 HCV patient samples with resistance-associated variants (RAVs) observed (Top), and the percentage of patients that had given RAVs as majority nucleotide (‘fixed sample’) (Bottom)**.

**S8 Fig. Comparison of simulated and observed *in vivo* mutation frequencies of HCV from 195 patients**.

**S9 Fig. Comparisons between HCV mutation rates estimated from Gellers et al. (2016)’s *in vitro* dataset and our *in vivo* dataset using nonsense mutation frequencies (A)**. Correlation between estimated in vivo and in vitro mutation rates are shown in B.

**S10 Fig. Simulations results showing the relationship between number of samples, coverage, and accuracy of inference, as determined by the correlation coefficient between simulated fitness costs and their estimates based on a mean-field approach**.

**S11 Fig. Diagram showing our approach to observe *in vivo* evolution of viruses. S12 Fig. Phylogenetic tree of HCV 1a used in this study**.

**S1 Table. Results of statistical tests comparing synonymous and non-synonymous mutation frequencies within each gene**.

**S2 Table. List of features included in random forest classification and regression models run on mutation frequencies and selection coefficients**.

**S3 Table. The results of beta regression with the estimated effects of different factors on transition mutation frequencies**.

**S4 Table. The results of beta regression with the estimated effects of different factors on inferred selection coefficients**.

**S5 Table. Resistance-associated variants in the NS3, NS5A, and NS5B genes examined in our study**.

**S1 Text (SI File). Validation of our frequency based methods: Between-host variability is consistent with within-host variability in HCV**.

**S2 Text (SI File). Supplementary Methods: Validating the estimated mutation rates**.

## Data availability

Processed bam files of HCV from each patient are available at https://figshare.com/articles/dataset/HCV_Bam_Files/13239491?file=25498772. All other relevant data are within the manuscript and its Supporting Information files.

## References

1. Bush RM, Bender CA, Subbarao K, Cox NJ, Fitch WM. Predicting the Evolution of Human Influenza A. Science. 1999;286: 1921–1925.

2. łuksza M, Lässig M. A predictive fitness model for influenza. Nature. 2014;507: 57–61. doi:10.1038/nature13087

3. Gaunt E, Wise HM, Zhang H, Lee LN, Atkinson NJ, Nicol MQ, et al. Elevation of CpG frequencies in influenza A genome attenuates pathogenicity but enhances host response to infection. Elife. 2016;5: e12735. doi:10.7554/eLife.12735

4. Crotty S, Cameron CE, Andino R. RNA virus error catastrophe: Direct molecular test by using ribavirin. Proceedings of the National Academy of Sciences. 2001;98: 6895–6900. doi:10.1073/pnas.111085598

5. Sanjuan R, Nebot MR, Chirico N, Mansky LM, Belshaw R. Viral Mutation Rates. Journal of Virology. 2010;84: 9733–9748. doi:10.1128/JVI.00694-10

6. Acevedo A, Brodsky L, Andino R. Mutational and fitness landscapes of an RNA virus revealed through population sequencing. 2014; 686–90. doi:http://dx.doi.org/10.1038/nature12861

7. Sanjuán R, Domingo-Calap P. Mechanisms of viral mutation. Cell Mol Life Sci. 2016;73: 4433–4448. doi:10.1007/s00018-016-2299-6

8. Trouplin V, Salvatori F, Cappello F, Obry V, Brelot A, Heveker N, et al. Determination of coreceptor usage of human immunodeficiency virus type 1 from patient plasma samples by using a recombinant phenotypic assay. J Virol. 2001;75: 251–259. doi:10.1128/JVI.75.1.251-259.2001

9. Theys K, Feder AF, Gelbart M, Hartl M, Stern A, Pennings PS. Within-patient mutation frequencies reveal fitness costs of CpG dinucleotides and drastic amino acid changes in HIV. Bloom JD, editor. PLoS Genet. 2018;14: e1007420–24. doi:10.1371/journal.pgen.1007420

10. Pennings PS, Kryazhimskiy S, Wakeley J. Loss and Recovery of Genetic Diversity in Adapting Populations of HIV. Fraser C, editor. PLoS Genet. 2014;10: e1004000–9. doi:10.1371/journal.pgen.1004000

11. Zanini F, Puller V, Brodin J, Albert J, Neher RA. In vivo mutation rates and the landscape of fitness costs of HIV-1. Virus Evolution. 2017;3: 10312.–12. doi:10.1093/ve/vex003

12. Kautz T, Forrester N. RNA Virus Fidelity Mutants: A Useful Tool for Evolutionary Biology or a Complex Challenge? Viruses. 2018;10: 600–17. doi:10.3390/v10110600

13. Cuevas JM, González-Candelas F, Moya A, Sanjuán R. Effect of Ribavirin on the Mutation Rate and Spectrum of Hepatitis C Virus In Vivo. Journal of Virology. 2009;83: 5760–5764. doi:10.1128/JVI.00201-09

14. Ribeiro RM, Li H, Wang S, Stoddard MB, Learn GH, Korber BT, et al. Quantifying the Diversification of Hepatitis C Virus (HCV) during Primary Infection: Estimates of the In Vivo Mutation Rate. Wilke CO, editor. PLoS Pathog. 2012;8: e1002881–13. doi:10.1371/journal.ppat.1002881

15. Geller R, Estada ú, Peris JB, Andreu I, Bou J-V, Garijo R, et al. Highly heterogeneous mutation rates in the hepatitis C virus genome. 2016;1: 16045. doi:10.1038/nmicrobiol.2016.45

16. Cheng X, Virk N, Chen W, Ji S, Ji S, Sun Y, et al. CpG usage in RNA viruses: data and hypotheses. Burk RD, editor. PLoS ONE. 2013;8: e74109. doi:10.1371/journal.pone.0074109

17. Caudill VR, Qin S, Winstead R, Kaur J, Tisthammer K, Pineda EG, et al. CpG-creating mutations are costly in many human viruses. Evol Ecol. 2020;34: 339–359. doi:10.1007/s10682-020-10039-z

18. Kato N. Genome of Human Hepatitis C Virus (HCV): Gene organization, sequence diversity, and variation. Microbial Comparative Genomics. 2000;5: 129–151.

19. Gray RR, Parker J, Lemey P, Salemi M, Katzourakis A, Pybus OG. The mode and tempo of hepatitis C virus evolution within and among hosts. BMC Evolutionary Biology. 2011;11: 113. doi:10.1186/1471-2148-11-131

20. Mauger DM, Golden M, Yamane D, Williford S, Lemon SM, Martin DP, et al. Functionally conserved architecture of hepatitis C virus RNA genomes. Proc Natl Acad Sci USA. 2015;112: 3962–3697. doi:10.1073/pnas.1416266112

21. Poveda E, Wyles DL, Mena á, Pedreira JD, Castro-Iglesias á, Cachay E. Update on hepatitis C virus resistance to direct-acting antiviral agents. Antiviral Research. 2014;108: 181–191. doi:10.1016/j.antiviral.2014.05.015

22. Lontok E, Harrington P, Howe A, Kieffer T, Lennerstrand J, Lenz O, et al. Hepatitis C virus drug resistance-associated substitutions: State of the art summary. Hepatology. 2015;62: 1623–1632. doi:10.1002/hep.27934

23. Sarrazin C. The importance of resistance to direct antiviral drugs in HCV infection in clinical practice. 2015; 1–19. doi:10.1016/j.jhep.2015.09.011

24. Kim S, Han K-H, Ahn SH. Hepatitis C Virus and Antiviral Drug Resistance. Gut and Liver. 2016;10: 890–895. doi:10.5009/gnl15573

25. Wyles DL, Luetkemeyer AF. Understanding Hepatitis C Virus Drug Resistance: Clinical Implications for Current and Future Regimens. Top Antivir Med. 2017;25: 103–109.

26. Sorbo MC, Cento V, Di Maio VC, Howe AYM, Garcia F, Perno CF, et al. Hepatitis C virus drug resistance associated substitutions and their clinical relevance: Update 2018. Drug Resistance Updates. 2018;37: 17–39. doi:10.1016/j.drup.2018.01.004

27. Baird NA, Etter PD, Atwood TS, Currey MC, Shiver AL, Lewis ZA, et al. Rapid SNP Discovery and Genetic Mapping Using Sequenced RAD Markers. Fay JC, editor. PLoS ONE. 2008;3: e3376. doi:10.1371/journal.pone.0003376.g003

28. Li JZ, Paredes R, Ribaudo HJ, Svarovskaia ES, Metzner KJ, Kozal MJ, et al. Low-frequency HIV-1 drug resistance mutations and risk of NNRTI-based antiretroviral treatment failure: a systematic review and pooled analysis. JAMA. 2011;305: 1327–1335. doi:10.1001/jama.2011.375

29. Echeverría N, Betancour G, Gámbaro F, Hernández N, López P, Chiodi D, et al. Naturally occurring NS3 resistance-associated variants in hepatitis C virus genotype 1: Their relevance for developing countries. Virus Research. 2016;223: 140–146. doi:10.1016/j.virusres.2016.07.008

30. Keck Z, Girard-Blanc C, Wang W, Lau P, Zuiani A, Rey FA, et al. Antibody Response to Hypervariable Region 1 Interferes with Broadly Neutralizing Antibodies to Hepatitis C Virus. Ou Jhj, editor. Journal of Virology. 2016;90: 3112–3122. doi:10.1128/JVI.02458-15

31. Perales C, Lorenzo-Redondo R, López-Galíndez C, Martínez MA, Domingo E. Mutant spectra in virus behavior. Future Virology. 2010;5: 679–698. doi:10.2217/fvl.10.61

32. Quan Y, Xu H, Wainberg MA. Defective HIV-1 quasispecies in the form of multiply drug-resistant proviral DNA within cells can be rescued by superinfection with different subtype variants of HIV-1 and by HIV-2 and SIV. J Antimicrob Chemother. 2014;69: 21–27. doi:10.1093/jac/dkt326

33. Díaz-Muñoz SL, Sanjuán R, West S. Sociovirology: Conflict, Cooperation, and Communication among Viruses. Cell Host Microbe. 2017;22: 437–441. doi:10.1016/j.chom.2017.09.012

34. Vignuzzi M, López CB. Defective viral genomes are key drivers of the virus-host interaction. Nat Microbiol. 2019;4: 1075–1087. doi:10.1038/s41564-019-0465-y

35. Andino R, Domingo E. Viral quasispecies. Virology. 2015;479–480: 46–51. doi:10.1016/j.virol.2015.03.022

36. Domingo E, Perales C. Viral quasispecies. PLoS Genet. 2019;15: e1008271. doi:10.1371/journal.pgen.1008271

37. McCloskey RM, Liang RH, Joy JB, Krajden M, Montaner JSG, Harrigan PR, et al. Global Origin and Transmission of Hepatitis C Virus Nonstructural Protein 3 Q80K Polymorphism. The Journal of Infectious Diseases. 2014;211: 1288–1295. doi:10.1093/infdis/jiu613

38. Echeverría N. Hepatitis C virus genetic variability and evolution. WJH. 2015;7: 831–16. doi:10.4254/wjh.v7.i6.831

39. Klein MB, Saeed S, Yang H, Cohen J, Conway B, Cooper C, et al. Cohort profile: the Canadian HIV-hepatitis C co-infection cohort study. Int J Epidemiol. 2010;39: 1162–1169. doi:10.1093/ije/dyp297

40. Bushnell B. BBTools. 2018.

41. Li H, Durbin R. Fast and accurate short read alignment with Burrows-Wheeler transform. Bioinformatics. 2009;25: 1754–1760. doi:10.1093/bioinformatics/btp324

42. R Core Team. R: A Language and Environment for Statistical Computing. Vienna, Austria; 2021.

43. McCarthy CJ, Whittaker TA, Boyle LH, Eyal M. Quantitative Approaches to Group Research: Suggestions for Best Practices. J SPEC GROUP WORK. 2017;42: 3–16. doi:10.1080/01933922.2016.1264520

44. Fox EJ, Reid-Bayliss KS, Emond MJ, Loeb LA. Accuracy of Next Generation Sequencing Platforms. Next Gener Seq Appl. 2014;1: 1000106. doi:10.4172/jngsa.1000106

45. Tisthammer KH, Kline C, Rutledge T, Diedrich CR, Ita S, Lin PL, et al. SIV evolutionary dynamics in cynomolgus macaques during SIV-Mycobacterium tuberculosis co-infection. Viruses. 2021;In press.

46. Wright S. Evolution in Mendelian Populations. Genetics. 1931;16: 97–159.

47. Haldane J. The effect of variation on fitness. The American Naturalist. 1937;71: 337–349.

48. Hartl DL, Clark AG. Principles of population genetics. Third. Sunderland, MA: Sinauer Associates, Inc; 1997.

49. Cribari-Neto F, Zeileis A. Beta Regression in R. Journal of Statistical Software. 2010;34: 1–24. doi:10.18637/jss.v034.i02

50. Ypma J, Borchers HW, Eddelbuettel D. Package ‘nloptr.’ 2018 Oct pp. 1–46. Available: https://cran.r-project.org/web/packages/nloptr/nloptr.pdf

51. Pedregosa F, Varoquaux G, Gramfort A, Michel V, Thirion B, Grisel O, et al. Scikit-learn: Machine Learning in Python. Journal of Machine Learning Research. 2011;12: 2825–2830.

