## Supplementary material for "Assessing *in vivo* mutation frequencies and creating a high-resolution genome-wide map of fitness costs of Hepatitis C virus": SupportingInfo.Table.Fig.pdf

**S1 Table. Results of statistical tests comparing synonymous and non-synonymous mutation frequencies within each gene.** Mann-Whiteny test was used with the Holm correction for multiple comparisons.

| Gene | corrected *P*-value |
|---|---|
| Core | 7.60E-26 |
| E1 | 3.03E-41 |
| HVR1 | 0.087466 |
| E2 | 1.36E-90 |
| NS1(P7) | 1.24E-11 |
| NS2 | 9.42E-47 |
| NS3 | 2.65E-184 |
| NS4A | 2.01E-13 |
| NS4B | 4.27E-74 |
| NS5A | 5.25E-89 |
| NS5B | 1.12E-40 |

**S2 Table. List of features included in random forest classification and regression models run on mutation frequencies and selection coefficients.** AA: amino acid, refN: ancestral nucleotide, bigAAChange: mutations resulting in a drastic amino acid change, MutAA: resulting amino acid from the mutation, ogAA: ancestral amino acid.

| | Features used in random forest estimations | | | | |
|---|---|---|---|---|---|
| 1 | bigAAChange | 29 | H-MutAA | 57 | Q-ogAA |
| 2 | 5' UTR | 30 | I-MutAA | 58 | R-ogAA |
| 3 | Core | 31 | K-MutAA | 59 | S-ogAA |
| 4 | E1 | 32 | L-MutAA | 60 | T-ogAA |
| 5 | E2 | 33 | M-MutAA | 61 | V-ogAA |
| 6 | HVR1 | 34 | N-MutAA | 62 | W-ogAA |
| 7 | NS1 | 35 | P-MutAA | 63 | Y-ogAA |
| 8 | NS2 | 36 | Q-MutAA | 64 | a-refN |
| 9 | NS3 | 37 | R-MutAA | 65 | c-refN |
| 10 | NS4A | 38 | S-MutAA | 66 | g-refN |
| 11 | NS4B | 39 | T-MutAA | 67 | t-refN |
| 12 | NS5A | 40 | V-MutAA | 68 | makesCpG |
| 13 | NS5B | 41 | W-MutAA | 69 | makesApA |
| 14 | RNAstructure | 42 | Y-MutAA | 70 | makesApC |
| 15 | Nonsyn | 43 | *-ogAA | 71 | makesApG |
| 16 | Positive AA | 44 | A-ogAA | 72 | makesApT |
| 17 | Negative AA | 45 | C-ogAA | 73 | makesCpA |
| 18 | Hydrophobic AA | 46 | D-ogAA | 74 | makesCpC |
| 19 | Polar AA | 47 | E-ogAA | 75 | makesCpT |
| 20 | Nonpolar AA | 48 | F-ogAA | 76 | makesGpA |
| 21 | Acidic AA | 49 | G-ogAA | 77 | makesGpC |
| 22 | *-MutAA | 50 | H-ogAA | 78 | makesGpG |
| 23 | A-MutAA | 51 | I-ogAA | 79 | makesGpT |
| 24 | C-MutAA | 52 | K-ogAA | 80 | makesTpA |
| 25 | D-MutAA | 53 | L-ogAA | 81 | makesTpC |
| 26 | E-MutAA | 54 | M-ogAA | 82 | makesTpG |
| 27 | F-MutAA | 55 | N-ogAA | 83 | makesTpT |
| 28 | G-MutAA | 56 | P-ogAA | | |

**S3 Table. The results of beta regression with the estimated effects of different factors on transition mutation frequencies.** The intercept is estimated for synonymous, non CpG creating A→G mutations. The results are from the best fit model selected using the AIC scores. Effect size represents the estimated increase or decrease in mutation frequencies compared to the intercept.

| | Estimate | Z-value | *P*-value | Effects |
|---|---|---|---|---|
| Intercept | -4.3767 | -243.55 | 0 | |
| T | 0.1157 | 5.55 | <0.0001 | **12.3%** |
| C | -0.5323 | -25.96 | <0.0001 | **-41.3%** |
| G | -0.7222 | -31.47 | <0.0001 | **-51.4%** |
| CpG | -0.0725 | -5.39 | <0.0001 | **-7.0%** |
| Nonsyn | -0.7582 | -33.94 | <0.0001 | **-53.2%** |
| bigAA Change | -0.1249 | -8.89 | <0.0001 | **-11.7%** |
| Core | -0.2169 | -10.96 | <0.0001 | **-19.5%** |
| E1 | 0.0375 | 2.06 | <0.0001 | **3.8%** |
| HVR1 | 0.2357 | 5.09 | <0.0001 | **26.6%** |
| E2 | 0.0843 | 5.94 | <0.0001 | **8.8%** |
| NS1 | 0.0699 | 2.32 | <0.0001 | **7.2%** |
| NS2 | 0.0895 | 5.27 | <0.0001 | **9.4%** |
| NS4A | -0.0559 | -1.61 | 0.107 | -5.4% |
| NS5B | -0.1632 | -10.48 | <0.0001 | **-15.1%** |
| T:Nonsyn | -0.1861 | -6.94 | <0.0001 | **-17.0%** |
| C:Nonsyn | -0.1432 | -5.16 | <0.0001 | **-13.3%** |
| G:Nonsyn | 0.0930 | 3.24 | 0.0012 | **9.7%** |

**S4 Table. The results of beta regression with the estimated effects of different factors on inferred selection coefficients.** The same model as the best fit model for estimating mutation frequencies was used for comparison. See S1. Table legend for details. Effect size represents the estimated increase or decrease in selection coefficients compared to the intercept.

| | Estimate | Z-value | *P*-value | Effects |
|---|---|---|---|---|
| Intercept | -6.5497 | -242.851 | 0.0000 | |
| T | -0.2358 | -6.928 | <0.0001 | **-21.0%** |
| C | -0.5067 | -16.425 | <0.0001 | **-39.7%** |
| G | -0.5772 | -17.072 | <0.0001 | **-43.9%** |
| CpG | 0.0677 | 5.773 | <0.0001 | **7.0%** |
| Nonsyn | 0.7292 | 25.988 | <0.0001 | **107.3%** |
| bigAA Change | 0.1298 | 12.367 | <0.0001 | **13.9%** |
| Core | 0.0593 | 3.503 | 0.0005 | **6.1%** |
| E1 | -0.0878 | -4.933 | <0.0001 | **-8.4%** |
| HVR1 | -0.4570 | -7.670 | <0.0001 | **-36.7%** |
| E2 | -0.1040 | -7.284 | <0.0001 | **-9.9%** |
| NS1 | -0.1672 | -5.424 | <0.0001 | **-15.4%** |
| NS2 | -0.1346 | -7.725 | <0.0001 | **-12.6%** |
| NS4A | -0.0535 | -4.206 | <0.0001 | **-5.2%** |
| NS5B | 0.0558 | 4.094 | 0.0001 | **5.7%** |
| T:Nonsyn | 0.1685 | 4.666 | <0.0001 | **18.4%** |
| C:Nonsyn | 0.1716 | 5.185 | <0.0001 | **18.7%** |
| G:Nonsyn | -0.0891 | -2.474 | 0.0134 | **-8.5%** |

**S5 Table. Resistance-associated variants in the NS3, NS5A, and NS5B genes examined in our study.** AA: amino acid, NT: nucleotide, wt: wildtype (reference), res: resistant, Ts: transition mutation, Tv: transversion mutation

| *Variant* | *AA Position* | *wtAA* | *resAA* | *Gene* | *NT Position* | *Mutation Type* | *wtNT* | *resNT* |
|---|---|---|---|---|---|---|---|---|
| V36M | 36 | V | M | NS3 | 3543 | Ts | g | a |
| V36A | 36 | V | A | NS3 | 3544 | Ts | t | c |
| V36G | 36 | V | G | NS3 | 3544 | Tv | t | g |
| V36L | 36 | V | L | NS3 | 3543 | Tv | g | c |
| T54A | 54 | T | A | NS3 | 3597 | Ts | a | g |
| T54S | 54 | T | S | NS3 | 3598 | Tv | c | g |
| V55A | 55 | V | A | NS3 | 3601 | Ts | t | c |
| Y56H | 56 | Y | H | NS3 | 3603 | Ts | t | c |
| Q80K | 80 | Q | K | NS3 | 3675 | Tv | c | a |
| Q80R | 80 | Q | R | NS3 | 3676 | Ts | a | g |
| S122G | 122 | S | G | NS3 | 3801 | Ts | a | g |
| S122R | 122 | S | R | NS3 | 3803 | Tv | c | a |
| S122R | 122 | S | R | NS3 | 3803 | Tv | c | g |
| S122D | 122 | S | D | NS3 | 3801 | Ts | a | g |
| S122D | 122 | S | D | NS3 | 3802 | Ts | g | a |
| S122T | 122 | S | T | NS3 | 3802 | Tv | g | c |
| I132V | 132 | I | V | NS3 | 3831 | Ts | a | g |
| R155G | 155 | R | G | NS3 | 3900 | Ts | a | g |
| R155K | 155 | R | K | NS3 | 3901 | Ts | g | a |
| R155T | 155 | R | T | NS3 | 3901 | Tv | g | c |
| A156T | 156 | A | T | NS3 | 3903 | Ts | g | a |
| A156G | 156 | A | G | NS3 | 3904 | Tv | c | g |
| A156V | 156 | A | V | NS3 | 3904 | Ts | c | t |
| D168H | 168 | D | H | NS3 | 3939 | Tv | g | c |
| D168V | 168 | D | V | NS3 | 3940 | Tv | a | t |
| D168T | 168 | D | T | NS3 | 3939 | Ts | g | a |
| D168T | 168 | D | T | NS3 | 3940 | Tv | a | c |
| D168K | 168 | D | K | NS3 | 3939 | Ts | g | a |
| D168K | 168 | D | K | NS3 | 3941 | Tv | c | a |
| D168K | 168 | D | K | NS3 | 3941 | Tv | c | g |

| | | | | | | | | |
|---|---|---|---|---|---|---|---|---|
| D168L | 168 | D | L | NS3 | 3939 | Tv | g | c |
| D168L | 168 | D | L | NS3 | 3940 | Tv | a | t |
| D168R | 168 | D | R | NS3 | 3939 | Tv | g | c |
| D168R | 168 | D | R | NS3 | 3940 | Ts | a | g |
| D168A | 168 | D | A | NS3 | 3940 | Tv | a | c |
| D168E | 168 | D | E | NS3 | 3941 | Tv | c | a |
| D168E | 168 | D | E | NS3 | 3941 | Tv | c | g |
| D168G | 168 | D | G | NS3 | 3940 | Ts | a | g |
| D168Y | 168 | D | Y | NS3 | 3939 | Tv | g | t |
| I170T | 170 | I | T | NS3 | 3946 | Ts | t | c |
| L175M | 175 | L | M | NS3 | 3960 | Tv | c | a |
| L175M | 175 | L | M | NS3 | 3962 | Ts | a | g |
| K24R | 24 | K | R | NS5A | 6346 | Ts | a | g |
| K24N | 24 | K | N | NS5A | 6347 | Tv | g | c |
| K24N | 24 | K | N | NS5A | 6347 | Tv | g | t |
| M28A | 24 | M | A | NS5A | 6357 | Ts | a | g |
| M28A | 24 | M | A | NS5A | 6358 | Ts | t | c |
| M28V | 24 | M | V | NS5A | 6357 | Ts | a | g |
| M28T | 24 | M | T | NS5A | 6358 | Ts | t | c |
| M28G | 28 | M | G | NS5A | 6357 | Ts | a | g |
| M28G | 28 | M | G | NS5A | 6358 | Tv | t | g |
| Q30H | 30 | Q | H | NS5A | 6365 | Tv | a | c |
| Q30H | 30 | Q | H | NS5A | 6365 | Tv | a | t |
| Q30G | 30 | Q | G | NS5A | 6363 | Tv | c | g |
| Q30G | 30 | Q | G | NS5A | 6364 | Ts | a | g |
| Q30K | 30 | Q | K | NS5A | 6363 | Tv | c | a |
| Q30E | 30 | Q | E | NS5A | 6363 | Tv | c | g |
| Q30R | 30 | Q | R | NS5A | 6364 | Ts | a | g |
| L31F | 31 | L | F | NS5A | 6366 | Ts | c | t |
| L31F | 31 | L | F | NS5A | 6368 | Tv | g | c |
| L31F | 31 | L | F | NS5A | 6368 | Tv | g | t |
| L31M | 31 | L | M | NS5A | 6366 | Tv | c | a |
| L31I | 31 | L | I | NS5A | 6366 | Tv | c | a |
| L31I | 31 | L | I | NS5A | 6368 | Ts | g | a |
| L31I | 31 | L | I | NS5A | 6368 | Tv | g | c |
| L31I | 31 | L | I | NS5A | 6368 | Tv | g | t |

| | | | | | | | | |
|---|---|---|---|---|---|---|---|---|
| L31V | 31 | L | V | NS5A | 6366 | Tv | c | g |
| S38F | 38 | S | F | NS5A | 6388 | Ts | c | t |
| H58D | 58 | H | D | NS5A | 6447 | Tv | c | g |
| E62L | 62 | E | D | NS5A | 6459 | Tv | g | c |
| E62L | 62 | E | D | NS5A | 6460 | Tv | a | t |
| A92T | 92 | A | T | NS5A | 6549 | Ts | g | a |
| Y93H | 93 | Y | H | NS5A | 6552 | Ts | t | c |
| Y93N | 93 | Y | N | NS5A | 6552 | Tv | t | a |
| Y93R | 93 | Y | R | NS5A | 6552 | Ts | t | c |
| Y93R | 93 | Y | R | NS5A | 6553 | Ts | a | g |
| Y93C | 93 | Y | C | NS5A | 6553 | Ts | a | g |
| Y93S | 93 | Y | S | NS5A | 6553 | Tv | a | c |
| L159F | 159 | L | F | NS5B | 8109 | Ts | c | t |
| E237G | 237 | E | G | NS5B | 8344 | Ts | a | g |
| S282T | 282 | S | T | NS5B | 8479 | Tv | g | c |
| C316Y | 316 | C | Y | NS5B | 8581 | Ts | g | a |
| C316F | 315 | C | F | NS5B | 8581 | Tv | g | t |
| C316N | 316 | C | N | NS5B | 8580 | Tv | t | a |
| C316N | 316 | C | N | NS5B | 8581 | Ts | g | a |
| V321I | 321 | V | I | NS5B | 8595 | Ts | g | a |
| V321A | 321 | V | A | NS5B | 8596 | Ts | t | c |

**S1 Fig. Top 20 features identified as important from the random forest regression model for predicting *in vivo* HCV mutation frequencies.** See S2 Table for all features included in the model.

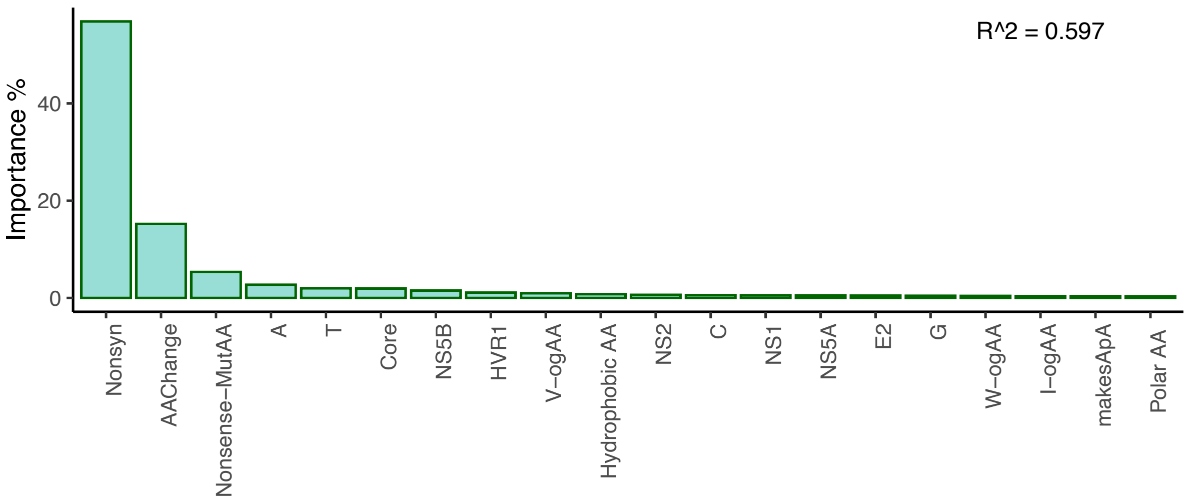

**S2 Fig. Comparison of *in vivo* and *in vitro* transition mutation frequencies in the HCV1a genome**. *In vivo* frequencies are from this study, and *in vitro* frequencies are from Geller et al. (2016). (A) Observed mutation frequencies by nucleotide, and (B) by gene.

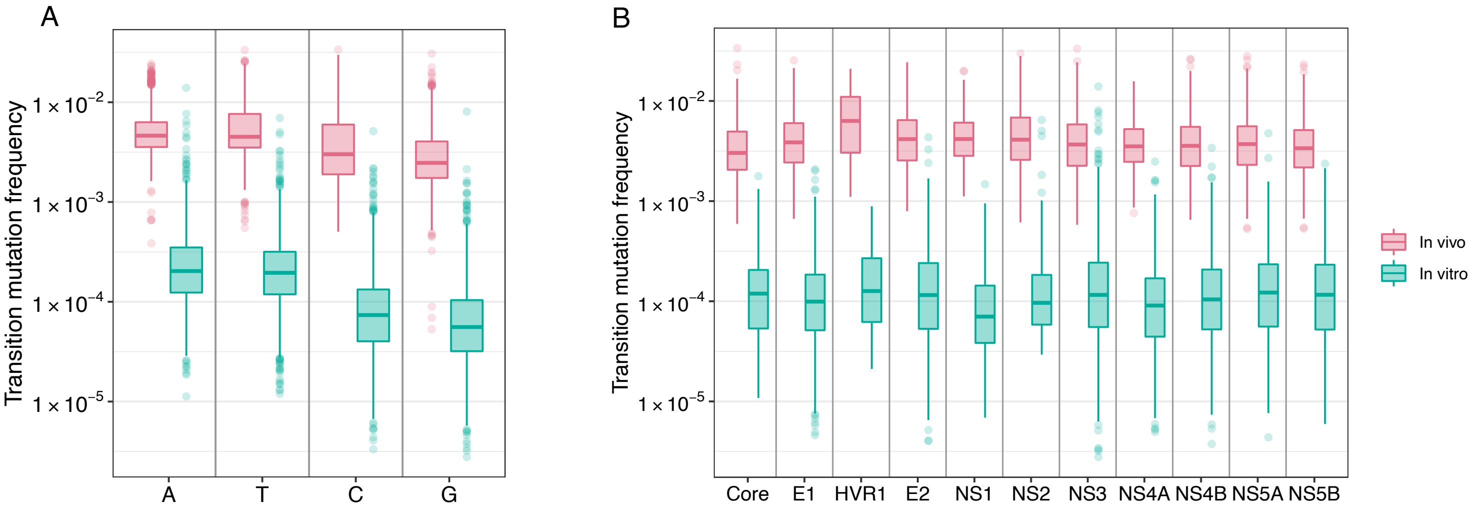

**S3 Fig. Transition mutation frequencies of the sites corresponding to the highly mutable (HM) and less mutable (LM) sites identified in Geller et al. (2016).** The dashed line represents the genome-wide mean mutation frequency in our 195 patient samples.

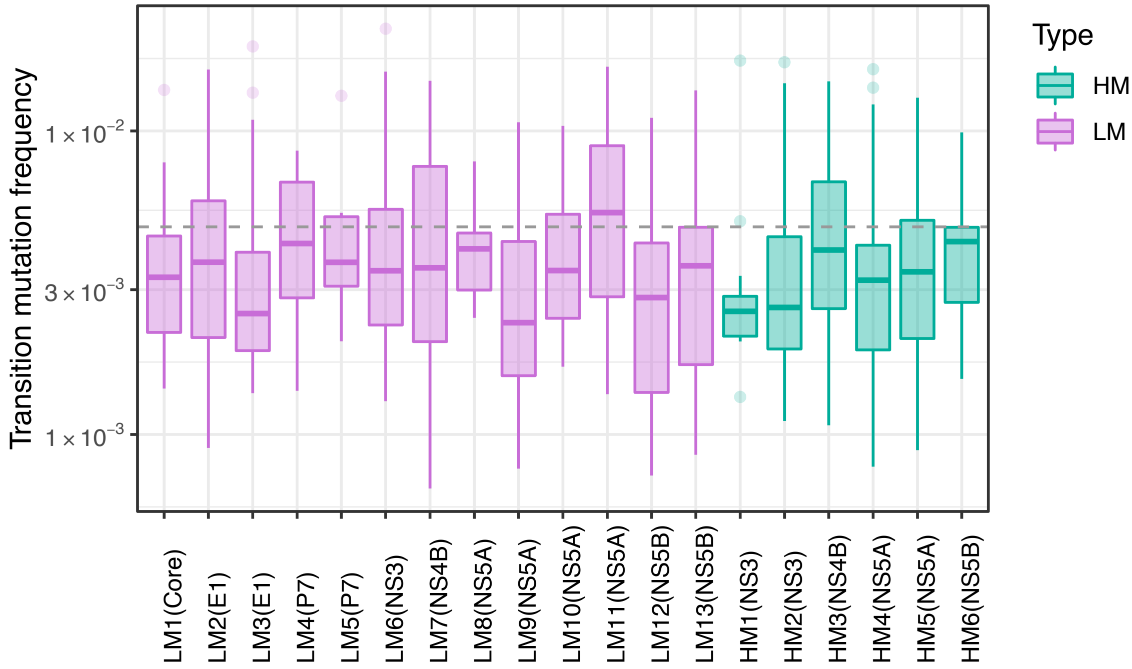

**S4 Fig. Mutation rates estimated from the dataset obtained from Geller et al. (2016).** Blue dots represent the estimated transition mutation rates for each nucleotide, and bars represent the 95% confidence intervals.

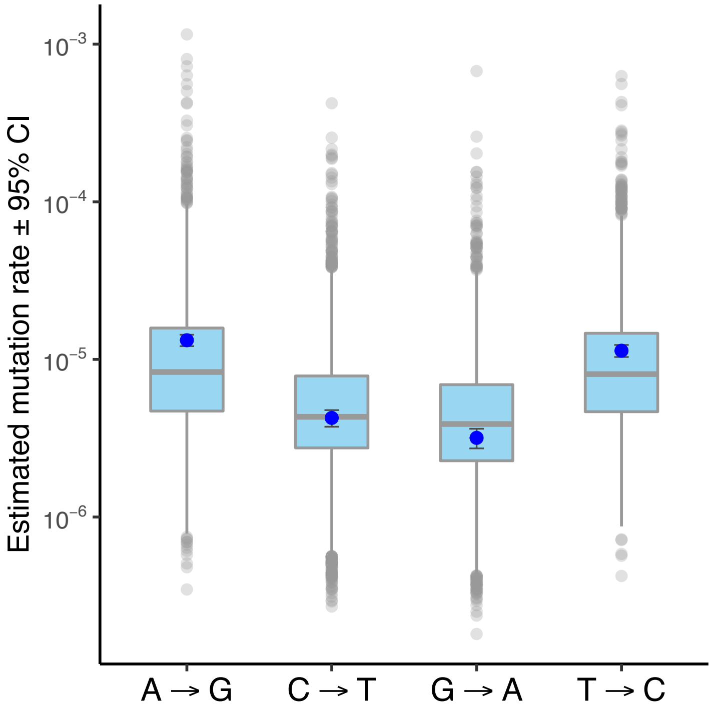

**S5 Fig. Average selection coefficients (A) and AT contents (B) of each genic region in the HCV genome.**

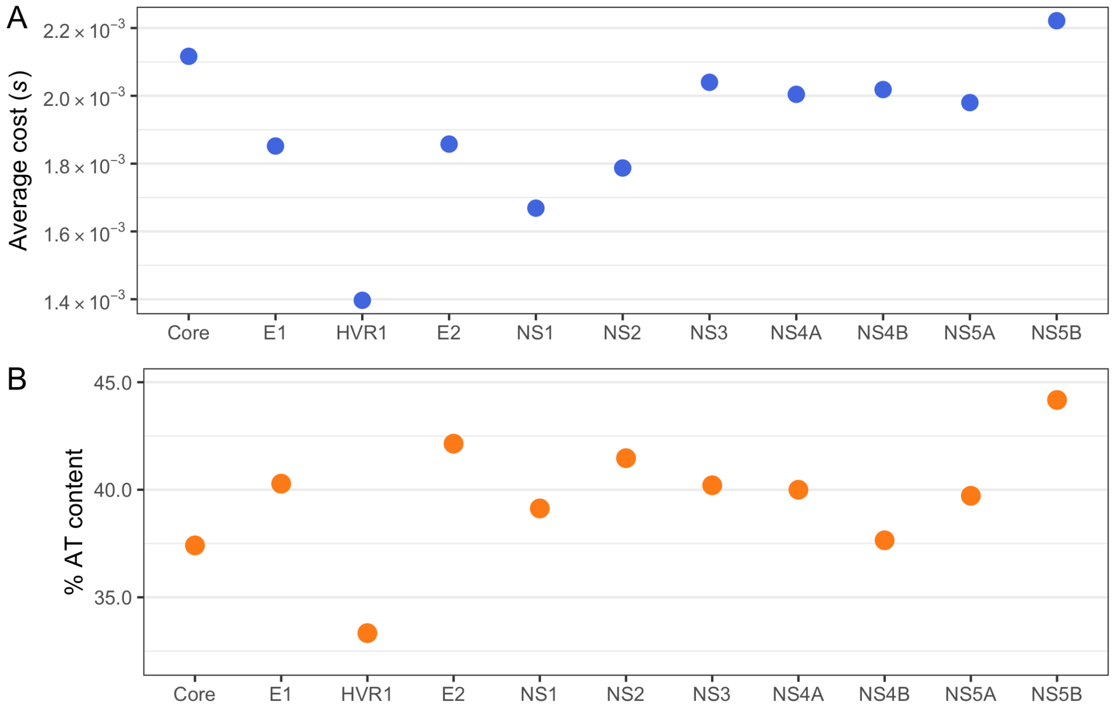

**S6 Fig. Distribution of fitness effects for each nucleotide (A, T, C, G) in the coding regions of the HCV genome, stratified by synonymous and nonsynonymous mutation status, using the estimated fitness costs (*s*) calculated in this study.**

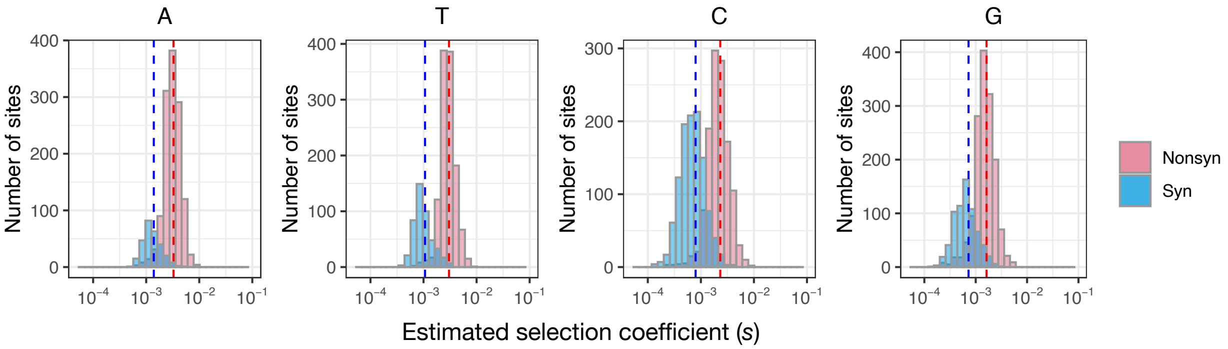

**S7 Fig. The percentage of HCV patient samples (n=195) with given resistance-associated variants (RAVs) (Top), and the percentage of patient samples that had given RAVs as majority nucleotide ('fixed sample') (Bottom).** X-axis shows all RAVs assessed in this study (S5 Table). Variant names in black are created by transition and those in brown are created by transversion mutations. Tv1 stands for transition mutations that result in C or A and Tv2 stands for transition mutations that results in G or T.

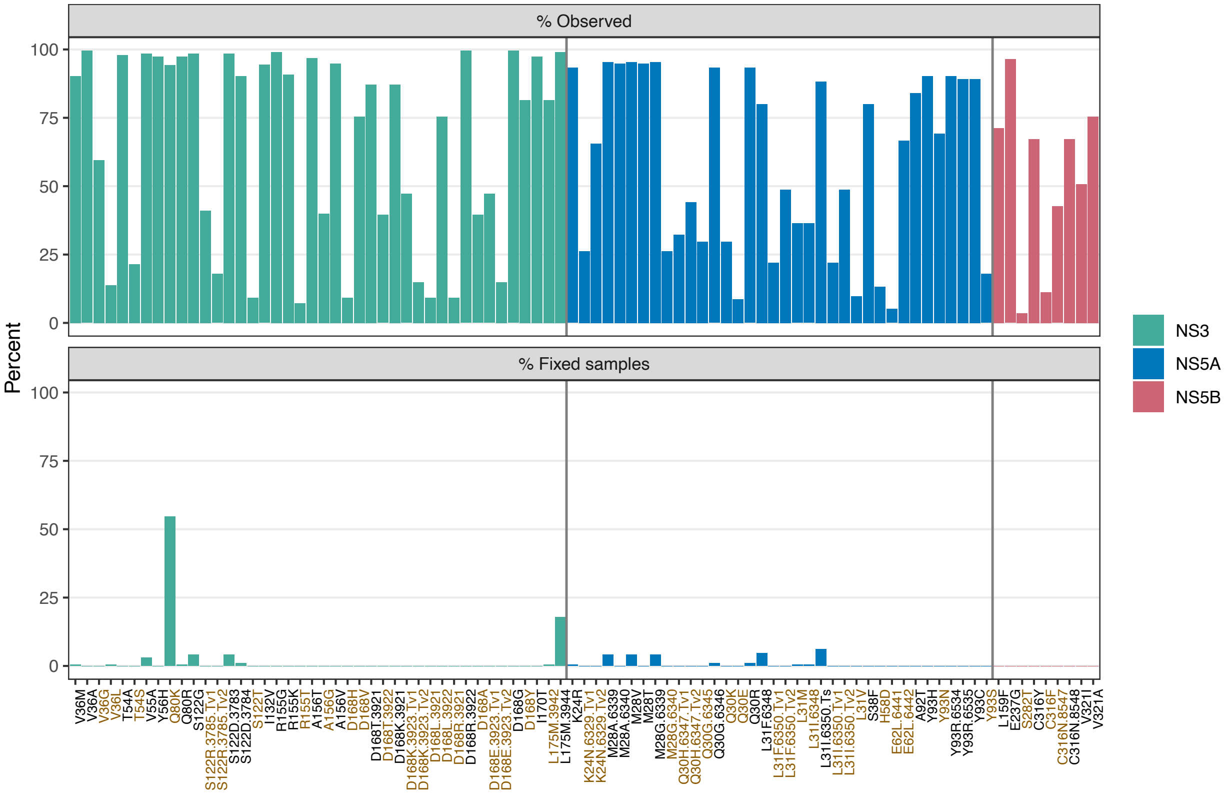

**S8 Fig. Comparison of simulated and observed *in vivo* mutation frequencies of HCV from 195 patients.** Examples from different types of mutations are shown. (A) Examples of sites that showed consistent patterns between simulated and observed frequencies (92.6 %) (adjusted P-value > 0.05), and (B) examples from the sites that showed different patterns (7.4 %) (adjusted P- values <0.05).

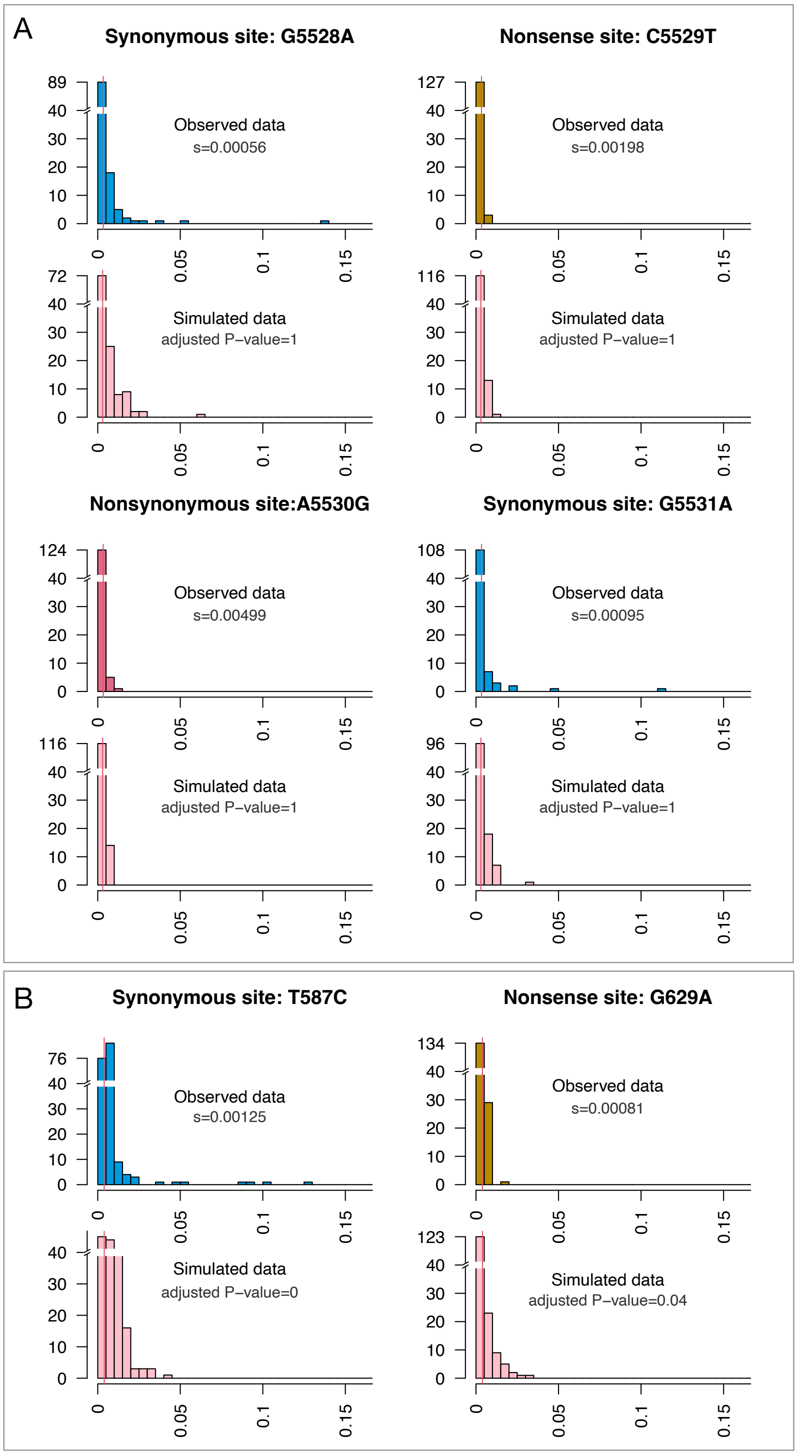

**S9 Fig. Comparisons between HCV mutation rates estimated from Gellers et al. (2016)'s *in vitro* dataset and our *in vivo* dataset using nonsense mutation frequencies (A).** Correlation between estimated *in vivo* and *in vitro* mutation rates are shown in B.

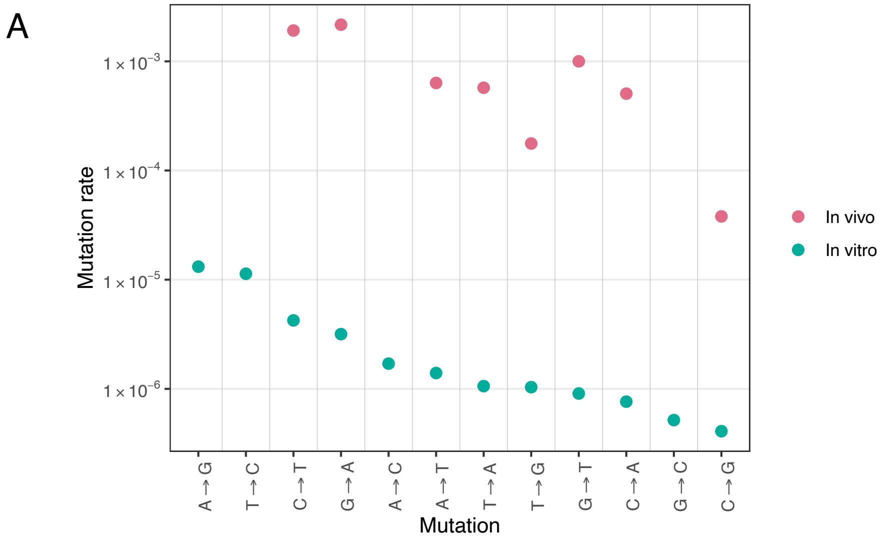


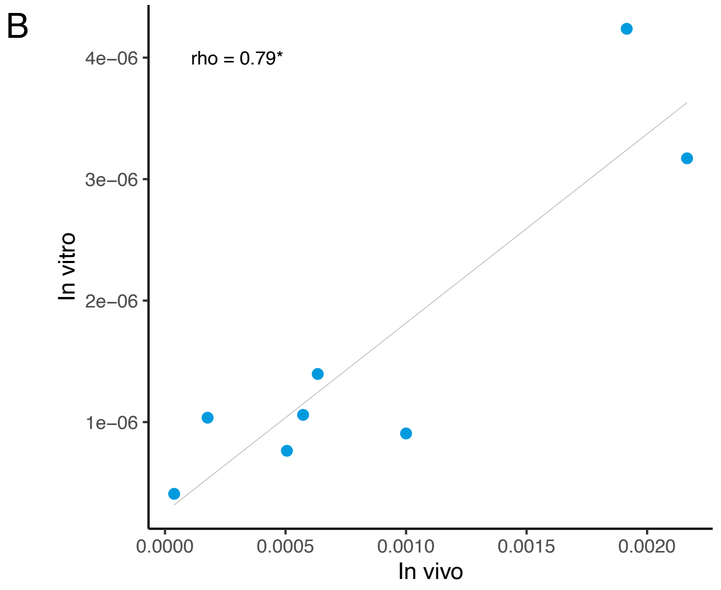

**S10 Fig. Simulations results showing the relationship between number of samples, coverage, and accuracy of inference, as determined by the correlation coefficient between simulated fitness costs and their estimates based on a mean-field approach.**

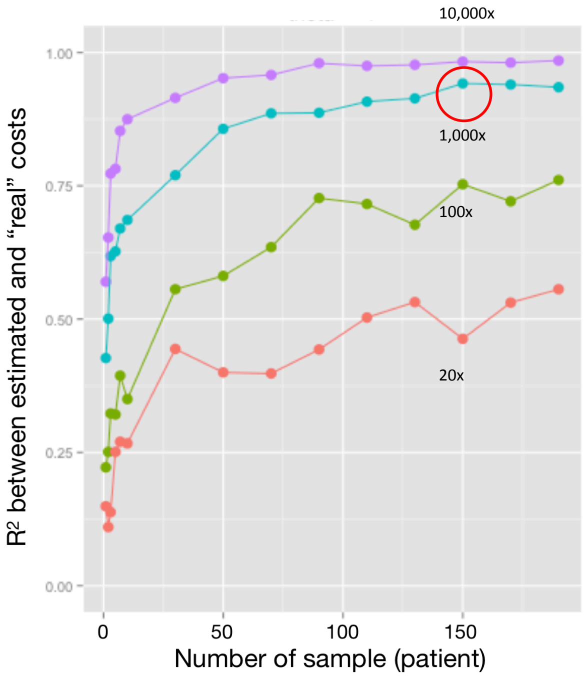

**S11 Fig. Diagram showing our approach to observe *in vivo* evolution of viruses.**

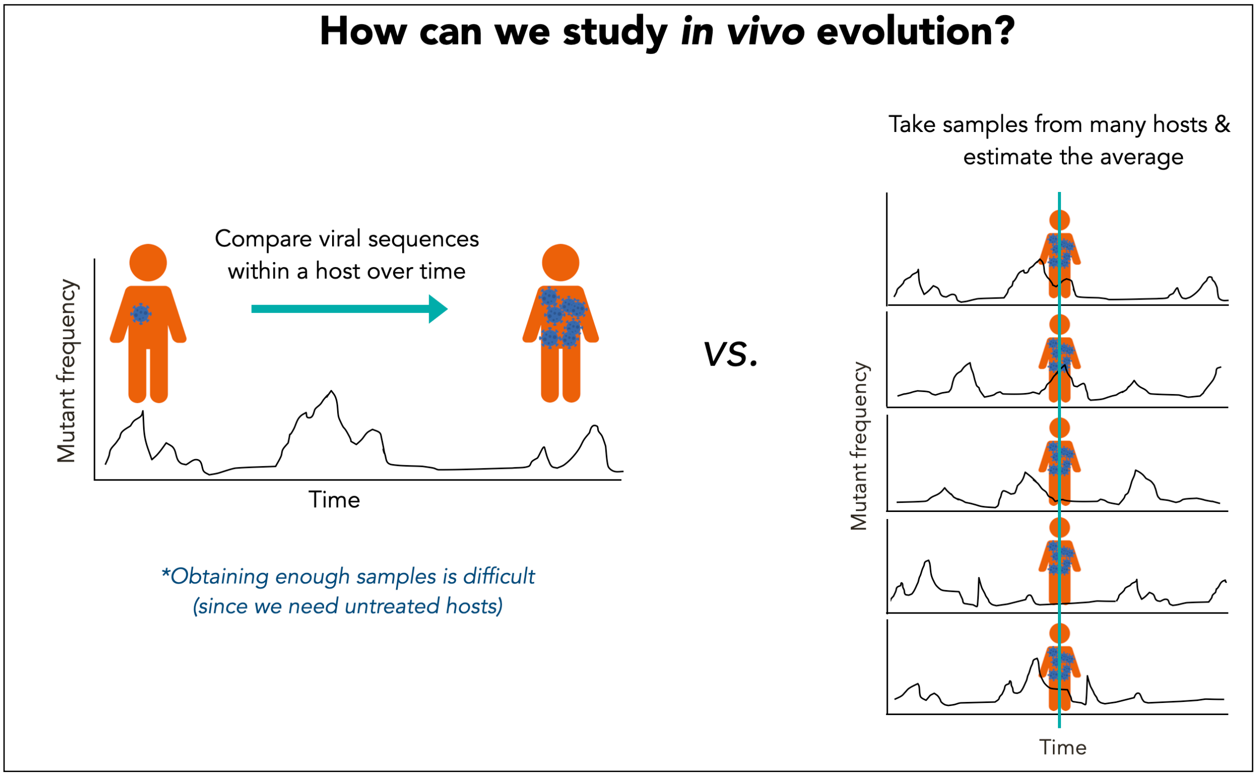

**S13 Fig. Phylogenetic tree of HCV 1a used in this study.** The numbers at the node represent the Bayesian posterior probability values.

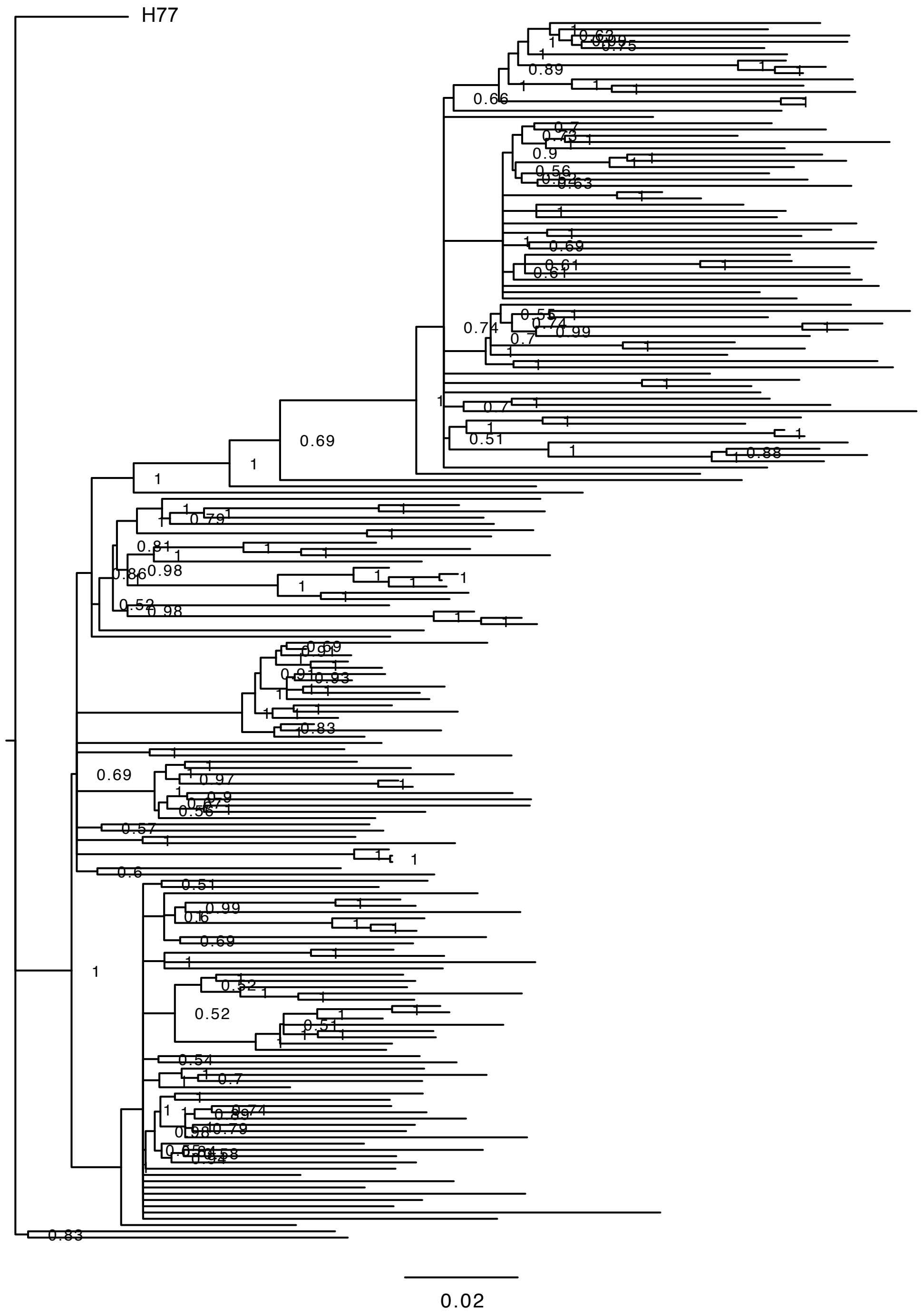

### Supporting Information (SI) File:

#### S1 Text: Validation of our frequency based methods: Between-host variability is consistent with within-host variability in HCV.

We assessed the relationship between genetic variability within hosts (*in vivo*) and between hosts by 1) testing the existence of correlation between the within-host and between-host nucleotide diversity at each nucleotide site (assuming that correlation should be 1), 2) checking the proportion of excluded sites through our filtering process to make sure that we did not eliminate too many sites/samples from our analysis to bias our results, and 3) assessing that the highly conserved sites between-host were also conserved within-host to further confirm that patterns of variability are consistent between-host and within-host.

First, we assessed nucleotide diversity levels between- and within-host using (1) average minor variant frequencies to represent within-host (*in vivo*) diversity and (2) Shannon's entropy indices of the HCV genome sequences downloaded from NCBI GenBank (n=423) along with consensus sequences of each viral population (*i.e.* each patient) from our dataset to represent between-host diversity. The two measures of diversity showed a high correlation with Spearman's $\rho = 0.691$ ($P < 2.2\text{x } 10^{-16}$) (SI File Fig 1), indicating that genome-wide evolutionary patterns were consistent between within-host and between-host. To put this correlation value ($\rho = 0.69$) into a context, we also calculated ρ values for comparisons between different subtypes. When within-host diversity of this study (HCV1a) was compared to between-host diversity of HCV subtype 1b (n=268) and 3a (n=540), ρ values decreased to 0.582 for subtype 1b and 0.527 subtype 3a, as expected (*i.e.* it was not a random association).

Next, we investigated potential effects of our filtering process, where we excluded the sites that had a majority nucleotide different from the reference nucleotide (H77) from each viral population (see Methods). If too many sites were excluded, and if these sites had high variability, this could mask within-host variability. We first calculated the proportion of excluded sites due to the nucleotide difference for each viral population, which was on average, only 6.6 (± 0.1) %. We also compared the consensus (majority nucleotide) sequences of all viral populations to gain further insights into the genetic variability between hosts. The overall sequence similarity (the average pairwise identity of consensus sequences) was relatively high (91.8%), and was within the range of the expected values (> 90% within a subtype) [1–3]. These results indicate that the number of sites excluded from the analysis was quite small, and the filtering step probably did not bias our results.

Lastly, we investigated if a group of highly conserved sites between-host also corresponded to highly conserved sites within-host. Alignments of all consensus sequences across the 195 genomes showed 49% of sites to be identical within the HCV coding regions. These identical sites, or highly conserved sites, occurred relatively evenly throughout the genome, except for in the HVR1 region, which had zero identical sites across the populations. Relative to non-identical sites, identical sites had a significantly lower (-5%) average mutation frequency ($4.69 \text{ x } 10^{-3}$ *vs*. $4.93 \text{ x } 10^{-3}$, Mann-Whitney test, $P = 0.0003$) and a significantly higher (+4%) average selection coefficient ($2.68 \text{ x } 10^{-3}$ *vs*. $2.58 \text{ x } 10^{-3}$, Mann-Whitney test, $P = 0.032$), confirming the correspondence between conserved sites between-host and within-host. Additionally, our recent study showed that close to 60% of sites in the HCV genome were conserved not only within subtypes but also between different genotypes/subtypes [4], indicating that the majority of sites are evolutionarily conserved (*i.e.* in mutation-selection balance).

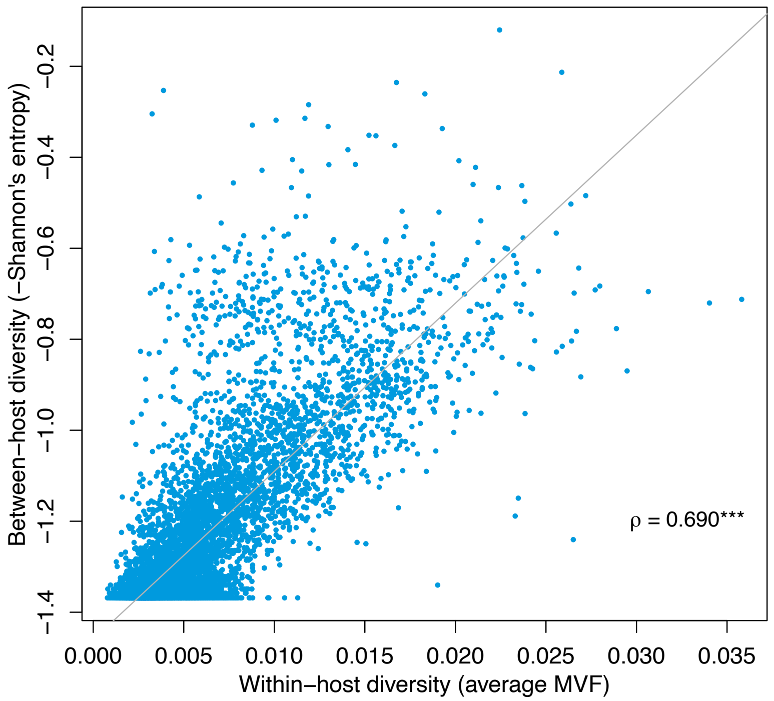


**S1 File Fig 1. Within-host (*in vivo*) *vs*. between-host nucleotide diversity comparison.** Between host diversity (Shannon's entropy) at each site was calculated from 618 HCV1a genome sequences (obtained from NCBI GenBank and the consensus sequences generated from each patient sample from this study). For within-host diversity, average minor variant frequencies (MVFs) of all samples at each site were used.

**S2 Text: Supplementary Methods: Validating the estimated mutation rates.**

The mutation rates estimated from Geller et al. [5] from the *in vitro* condition should not show the effects of different types of mutations, such as synonymous and nonsynonymous mutations. To validate this, we investigated the nucleotide-level transition mutation rates estimated from Geller et al. [1] using a beta-regression model, where the effects of different factors (types of mutations, location, RNA structure) on mutation rates were assessed. Since the dataset contained many zeros, a transformation [y = x (n − 1) + 0.5)/n ] (n = the sample size) were applied as recommended by [6]. The best fit model is shown below, which revealed no significant effects of nonsynonymous mutations, CpG creating mutations, and HVR1, as expected from the *in vitro* condition without selective pressures from the host's immune systems. The largest effects were due to the type of ancestral nucleotide, with the mutations rates of C→T and G→A being about a half of that of T→C and A→G. However, the model results show small effects of mutations that cause drastic amino acid change (bigAAChange) and locations (E1, NS1, NS2, NS5B), which we have no explanation for.

| | Estimate | Z-value | *P*-value | Effects |
|---|---|---|---|---|
| (Intercept) | -8.0072 | -512.50 | <0.0001 | 0% |
| t | -0.0438 | -2.471 | 0.0135 | -4.3% |
| c | -0.5938 | -32.163 | <0.0001 | -44.8% |
| g | -0.7159 | -37.205 | <0.0001 | -51.1% |
| bigAAChange | 0.0372 | 2.815 | 0.0049 | 3.8% |
| Core | -0.0604 | -1.991 | 0.0464 | -5.9% |
| E1 | -0.0737 | -2.6837 | 0.0073 | -7.1% |
| NS1 | -0.1961 | -3.9992 | 0.0001 | -17.8% |
| NS2 | -0.1938 | -7.1382 | <0.0001 | -17.6% |
| NS5B | -0.0596 | -3.5023 | 0.0005 | -5.8% |

To further assess the validity of the estimated mutation rates from Geller et al. [1], we estimated selection coefficients using mutation rates calculated at different levels; 1) an individual site-based level and 2) a quartile-based level, and compared to those estimated using a single, uniform mutation rate for each nucleotide used in the main analysis. The site-based mutation rates were calculated separately for every site along the genome. The quartile-based mutation rates were calculated by grouping the mutation rates into quartiles and averaging them. The results showed that the original method using a uniform mutation rate for each ancestral nucleotide produced the narrowest distributions of selection coefficients, compared to the other two methods for all four nucleotides, indicating the best approach. Examples from C→T mutations are shown in the figure below (SI File Fig 2.)

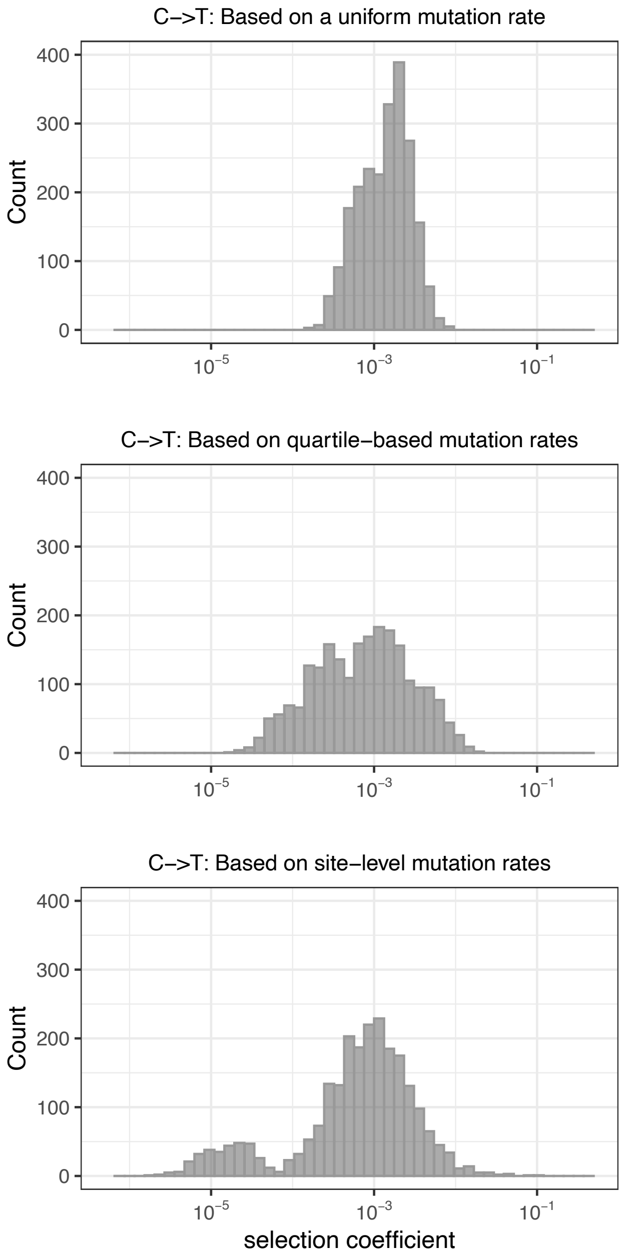


**SI File Fig 2. Estimated selection coefficients using different levels of mutation rates.**
